# Accuracy and Scalability of Machine Learning Methods for Genotype-Phenotype Association Data

**DOI:** 10.1101/2025.02.13.638022

**Authors:** Kieran Collienne, Lilin Zhang, Nicholas Sumpter, Meagan Leask, Michael Witbrock, Alex Gavryushkin

## Abstract

Many machine learning methods can be applied to predicting phenotypes from genetic data. Which of these methods work best remains an open question, however. To answer this question, we propose to compare a variety of approaches’ ability to predict a simulated non-linear complex trait. Specifically, we evaluate these methods on their accuracy and scalability with respect to the amount training data available, the noise present in the data, the complexity of the simulated (trait) functions, and their ability to provide insight into the simulated trait. We then compare the best approach to state-of-the-art models in real data, predicting gout in the UK Biobank. We find that transformer encoders outperform all other methods in simulations, and perform comparably to the state-of-the-art with real data, with a promise to scale to significantly larger datasets.

---

Simple linear models often fail to fully explain heritable diseases [76]. While more powerful methods are prone to overfitting, in practice linear models also often outperform complex models when the number of variables *p* (genomic predictors, typically single nucleotide variants or SNVs) is significantly greater than the number of samples *n* [23] (typically the number of sequenced individual organisms). In some cases this can be overcome by using data augmentation or pre-training [19], however the ideal solution is to fit the model using more data. Exactly how much data is required for complex trait prediction remains a largely unanswered question and an area of extensive research efforts [1, 22, 27, 11].

To address this, we design a simulation to determine which of our models should be expected to work, and under what circumstances. Our simulation aims to answer the following questions: How well do these models cope with noisy data typical for molecular biology? How does performance scale with available training data? Can the same model infer both simple and complex non-linear trait functions?

We define a class of functions of varying complexity, including both a linear and non-linear component, and assume our target trait can be accurately approximated using some function from this class. To evaluate our models, we simulate functions from this class that aim to approximate the effects regulatory SNVs might have on a complex trait. This model is partially inspired by linear models of expression Quantitative Trait Loci (eQTLs) [54], but our main goal was to identify a biologically meaningful genotype-phenotype association mechanism under which the generated data cannot be approximated by a linear (or in fact additive) model to a level of accuracy acceptable in biomedical applications, so classical genome-wide association study (GWAS) techniques would result in unsatisfactory predictions in this case.

To ensure our simulation has a realistic distribution of SNVs, as well as realistic relationships between them, we use sequences from the GISAID hCoV-19 data set [16]. We treat the first entry as the reference sequence, and consider every nucleotide to be a potential SNV. We encode these SNVs as 0 if the value matches the reference sequence, 1 otherwise, and identical sequences are removed from the data. Finally, we then simulate binary phenotypes using functions of varying complexity and with varying levels of noise. Fitting this data is therefore a binary classification problem. While continuous (or otherwise non-binary) complex traits exist, binary polygenic traits, disease susceptibility for instance, are one of the simplest cases in which classic genome-wide association studies fail.

## 1 Simulation Design

Gene expression is simulated as a linear combination of the presence or absence of specific SNVs, where certain SNVs *v_i_* have an effect *β_i_* on the expression of some gene.

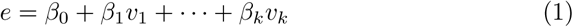

Binary gene effects *g* are present when *e* exceeds or is below a certain threshold. Since *β*_0_ is arbitrary we can say without loss of generality that effect *g* is present if and only if *e >* 0.

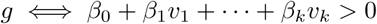

We say that a complex trait *T* is present if a certain logical function of effects *g*_1_*, …, g_n_* is true, e.g. *T* = (*g*_1_ ∧*g*_2_)∨¬*g*_3_. The complexity of the overall function has two dimensions, the number of genes *k* and the number of regulatory SNVs per gene *h*. We randomly sample a function *g* with complexity (*k, h*) as follows.

We start by randomly generating *k* gene regulatory functions *g*_1_*, …, g_k_*, each involving *h* SNVs. For each gene, we choose *h* regulating SNVs uniformly at random without replacement. Next, we choose *h* values 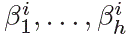 from *N* (0, 1). Finally, choose 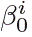 from *N* (0, 1). The function *g_i_* is then:

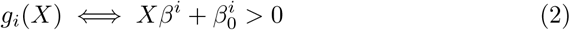

We combine these functions *g*_1_*, …, g_k_* into a logical formula represented by a decision tree with *k* leaves. This tree is constructed by recursively splitting the set of functions to form a tree (Fig. 1). We start with a set *S* = {*g*_1_*, …, g_h_*}, and a single node. This node is randomly assigned either ∧ or ∨, and two new branches, each containing half of *S*, are created. These steps are repeated recursively, halting when *S* contains a single gene function. Finally, we negate 10% of the leaves at random.

**Figure 1:**
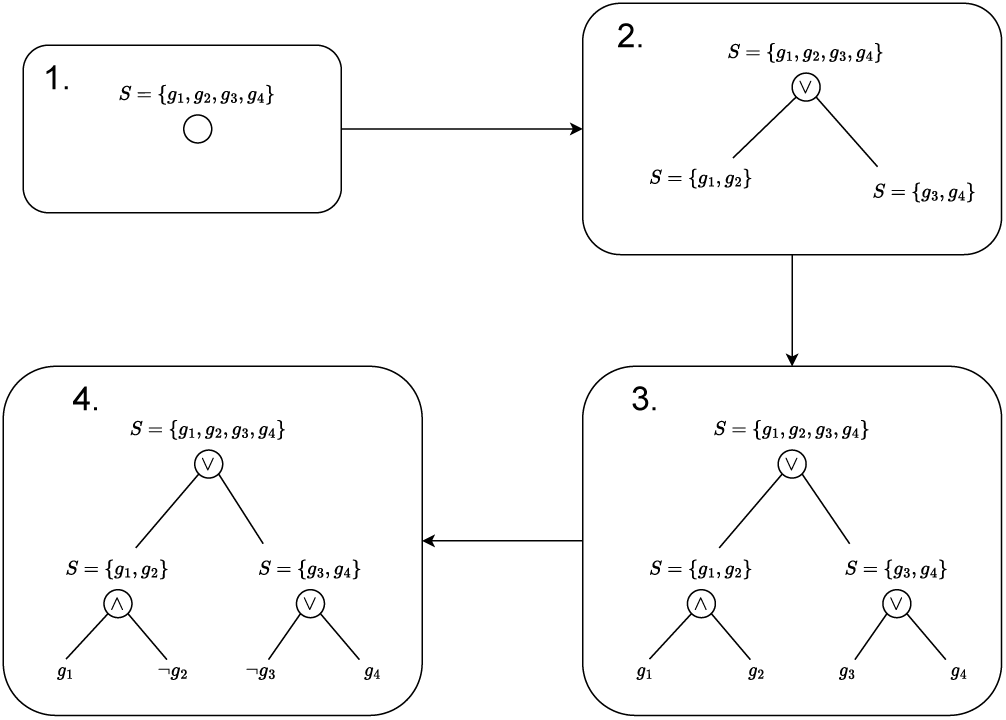
Decision tree sampling process. The final function is (*g*_1_ ∧ ¬*g*_2_) ∨ (¬*g*_3_ ∨ *g*_4_).

**Noise** We add noise to both the linear expression function *g* (Eq. (2)), and the output of the logical function (Fig. 1). In both cases we use a noise parameter as the probability of returning a random value instead of the true output. This random output in both cases is either 1 or 0, with a 50% chance of each.

We generate datasets with 2, 8, 32, or 128 SNVs per gene regulation function; 2, 8, 32, or 128 gene functions; and noise parameters of 0.0, 0.2, and 0.4. Note that we use the same noise parameter for both the eQTL gene regulatory functions and logical gene function. We reserve the final 30% of each dataset as a test set. The remaining data is either all used for training with hyperparameters chosen using cross-validation, or 10% is used as a validation set for hyperparameter selection, depending on the model. Note that rather than using all available training data, we use only a limited amount in each trial, to better understand how performance scales with different amounts of training data.

### 1.1 Models Tested

We fit the simulated data with several different models. In each case, we attempt to classify the output *o* as either having or not having the simulated trait, given the input SNV variants *v*_1_*, …, v_k_*.

#### 1.1.1 Logistic Regression

The simplest option, logistic regression, has often performed well in practice [8]. In this model we optimise parameters *β* to minimise the cross-entropy between the output *ô* ∈ (0, 1) and the simulated value *o*. Given *β*, the logistic regression function *lin* is:

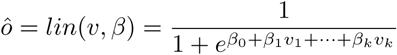

We implement logistic regression using PyTorch [49], fitting the parameters using AdamW [56].

#### 1.1.2 Gradient-Boosted Decision Trees

Instead of additive regression models, where each variant increases or decreases the risk of gout, we can use decision trees to directly predict gout based on the presence or absence of specific variants. Trees do not require SNV contributions to be additive, instead they represent logical functions of variants. If significant non-additive interactions are present, we expect tree-based approaches to find them more easily than regression. Instead of fitting a single tree, methods aggregating multiple trees often generalise better to unseen data than single trees [67]. Rather than randomly sampling subsets of features to train a range of trees, as in Tin Kam Ho [67], more recent approaches iteratively choose trees that minimise the current residual loss using gradient-boosting [21]. We therefore use LightGBM [26], a state-of-the art method for fitting gradient-boosted decision trees to large scale data.

While logistic regression cannot model the logical component of our simulation, decision trees cannot model the linear component. Nonetheless, we expect them to capture some component of the function, and we use these as a benchmark for other approaches.

#### 1.1.3 Support Vector Machines

For comparison we also include Linear Support Vector Classifiers, using the scikit-learn implementation [50]. We use hinge loss with the default l2 penalty, and choose the regularisation parameter *C* using 3-fold cross-validation over 32 trials. *C* is sampled from (10*^−^*^2^, 10^2^) on a log scale. Linear SVMs are only capable of perfectly classifying linearly separable data [14]. Since some logical functions are not linearly separable, we therefore expect SVMs to fail in some cases. Despite this, they may still provide a good approximation overall.

#### 1.1.4 Multi-Layer Perceptrons

The first of our models that is theoretically capable of perfectly modelling the simulation is the classic Multi-Layer Perceptron (MLP) [59]. The power of this model is determined by both the depth and width of the intermediate hidden layers, and we theoretically only need enough neurons in the first layer to represent each of our linear functions. The later layers need to approximate the logical component. Since a single-layer MLP can perform the and, or, and not operations, there is some arrangement of weights in which an MLP should perfectly capture the simulation.

#### 1.1.5 Transformer Encoder

We use the transformer encoder architecture shown in Fig. 2. SNVs are encoded using a learned embedding of each SNV token, where the tokens are either 0 or 1. We perform a separate hyperparameter search for each simulated dataset to avoid over– or under-fitting.

**Figure 2:**
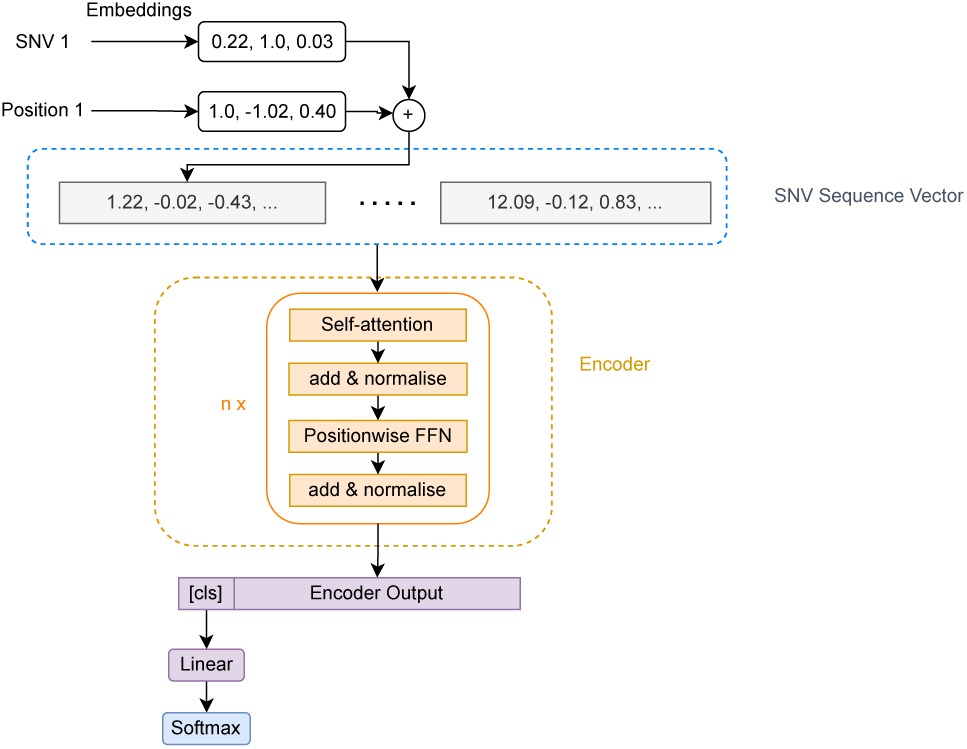
Transformer encoder used for simulations.

**Figure 3:**
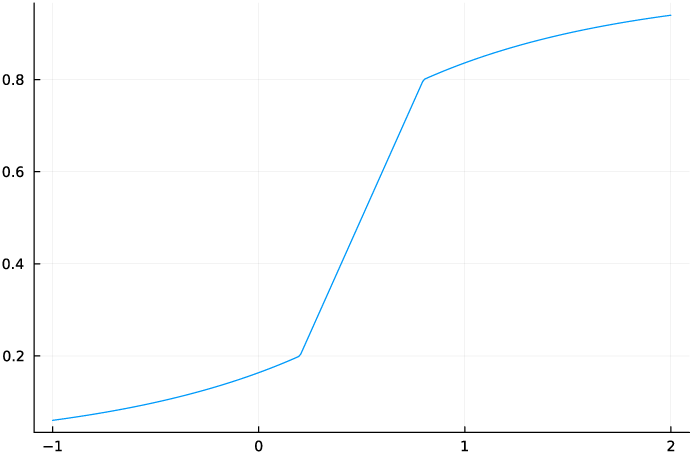
*y* = *r*(*x*) as defined in Eq. (11).

The hCoV-19 sequences we use are over 20, 000 variants long, and as a result transformers with the original quadratic self-attention mechanism take several hours to run a single training epoch. This in turn limits the number of hyperparameters that can be practically tested. We therefore include models with both the original scaled dot product attention mechanism and Linformer [72] for comparison.

## 2 Differentiable Logic

We designed a model to perfectly match the simulated function, using a combination of linear regression and differentiable fuzzy logic. This model shares our simulation’s assumption that the function we are approximating is a logical combination of over– or under-expressed genes, which are themselves modelled as a linear combination of SNVs.

### 2.1 Linear EQTLs

Following the simulation, our logical model supposes the expression of each gene *g_i_* is approximately 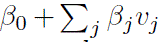, where *β_j_* is the contribution of the SNV *v_j_*. Furthermore, we assume that a complex trait *t* (gout) occurs when the expression of certain genes are above or below particular thresholds.

We propose a model that assumes the effects follow this pattern, without specifying the genes involved. Instead, we use implied expression values and optimise the entire model directly for complex trait prediction. Our model is based on a linear contribution of SNVs to expression of genes. Whether these expression levels exceed learned thresholds is used as a logical input to a disjunctive normal form expression, which is then used to predict the trait. We simultaneously optimise the disjunctive normal form and linear components.

### 2.2 Disjunctive Normal Form

A logical formula is said to be in disjunctive normal form if it is a disjunction of clauses, each of which is a conjunction of propositions or their negations.

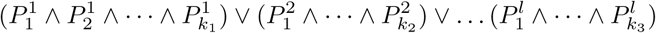

where ∨ is the logical ‘or’ function (disjunction), and ∧ is ‘and’ (conjunction). Every propositional formula is equivalent to a formula in disjunctive normal form [41]. For simplicity, we assume the logical function for each trait is in disjunctive normal form, e.g.

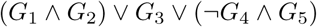

where *G_i_*is true if and only if *g_i_ > x_i_*.

We leave the length of this formula as an adjustable hyperparameter. We further assume that the genes we predict the expression *g_i_*of, are exactly the ones responsible for the trait *t*. Since we make no assumptions about which SNVs contribute to this gene, we can do this without loss of generality.

### 2.3 Model Definition

To simultaneously optimise both the logical and linear components of this model, we define the following differentiable logic, inspired by a many-valued logic defined by Łukasiewicz [34]. In Łukasiewicz logic, we have the following continuous definitions for the and, or, and not functions:

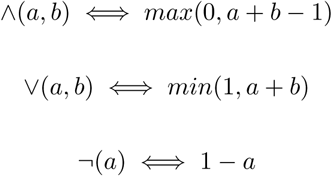

where *a*, and *b* are real numbers in the range [0, 1], each representing the confidence in some proposition. In our complex-trait model these are the variables *G_i_* indicating whether a gene is over or under expressed. Note that ∧ and ∨ are the definitions of strong (rather than weak) disjunction and conjunction. We do not use the implication, equivalence, weak conjunction or weak disjunction operations. Using a real-valued generalisation of classical logic allows us to differentiate the model and learn parameters with gradient-based methods. We need to be able to not only differentiate the correct logical function, but to choose what that function should be. For the latter, we introduce a *weighted* version of Łukasiewicz logic.

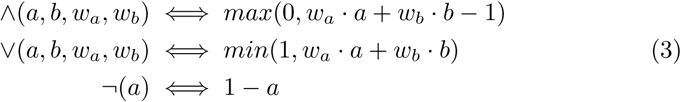

The weights *w_a_* and *w_b_* change the extent to which *a* and *b* are included in the ‘and’ or ‘or’ functions. Using these weights, we can extend these functions to many-valued functions by replacing *a* and *b* with an arbitrarily long vector *a*:

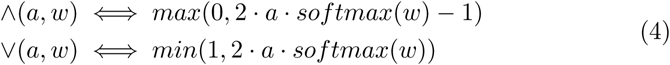

where *softmax*(*x*) is the function 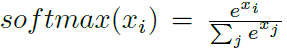. This allows us to use weights as a gradually adjustable form of variable selection. Finally, to ensure the function is differentiable outside the range [0, 1], we replace the *min*, and *max* hard thresholds with the function *r* (Eq. (11) and Fig. 17).

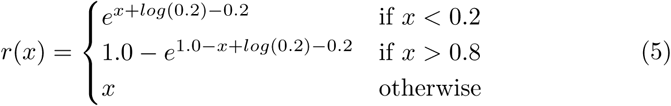

The final ‘and’ and ‘or’ functions are differentiable at all points, and outputs remain within the range (0, 1):

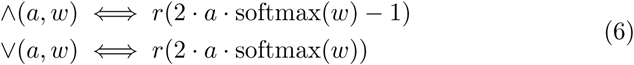

Using these operations, we define the model as follows. Expression values *e* are a linear combination of SNV values *X_snv_*, using weights 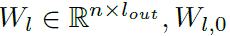, where *n* is the number of SNVs, and *l_out_* is a hyperparameter deciding how many genes are included in the model.

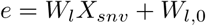

Next, we construct the matrix *X^′^* with each value and its negation *X^′^* = [*e,* ¬*e*]. We then get the values of the *c* conjunctions *a*_1_*, …, a_c_* weights 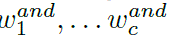:

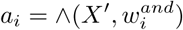

The final output is that of the disjunction, with variable selection weights *w^or^*:

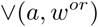

The complete model has learned parameters 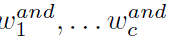 for each of the *c* conjunctions, *w^or^* for the disjunction, and *W_l_* for the linear function of SNVs. We implement our model using Lux [47] and MLJ [6], with the AdamW algorithm to optimise parameters [33].

**Binarised Model** When two weights *w_i_* and *w_j_* in *w* are much larger than all others, 2 · *softmax*(*w*) gives near zero weight to other inputs, while *i* and *j* will have weight near one if *w_i_* ≈ *w_j_*. With extreme weights, behaviour therefore approaches that of a binary model. This binary model has the classical logic ‘and’ and ‘or’ functions, and is easier to interpret than the weighted model. To convert a trained weighted model to a strictly binary one, we convert the weights from *w* ∈ R*^n^* to *w^′^* ∈ {0, 1}*^n^* as follows.

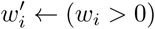

If 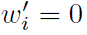 for all *i*, then we assign 1 to the index with the largest value: max*_i_ w_i_* ← 1. These are the weights that have increased, rather than decreased, which we interpret as meaning they should be included in the expression. Using these new weights *w^′^*, we revert to the hard [0, 1] limit functions in Eq. (9) and use the new weights directly without the softmax operation (Eq. (13)).

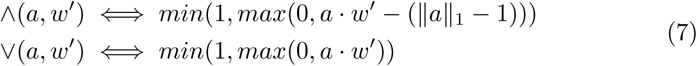

This allows us to learn parameters in the continuous model using gradient descent, then convert the learned continuous model to a classic binary model if the weights converge to extreme values.

## 3 Simulation Results

We evaluate the overall performance of each method using balanced accuracy, comparing each method’s performance with varying amounts of training data, levels of noise, and complexity of the simulated function. Note that the simulated data sets are not balanced, and up to 80% of samples may have the same class, which we address by using balanced accuracy rather than accuracy. We also include an ‘uninformed’ model, which always returns 1 or always returns 0, whichever is highest, as well as the true classification function in our analysis. Accuracy scores should be considered with respect to these values.

### 3.1 Overall Balanced Accuracy

After fitting every method to every simulated dataset, we calculate balanced accuracy for each. With identified true positives *TP*, true negatives *TN*, false positives *FP*, and false negatives *FN*, the balanced accuracy is:

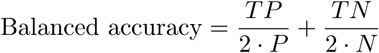

We additionally calculate the balanced accuracy for each dataset using an ‘uninformed’ method that always returns either all ones or all zeros, whichever scores highest. The mean balanced accuracy across all datasets with a particular training limit is shown in Fig. 4. Note the dashed line representing the balanced of the uninformed model. In several instances SVMs failed to train with small amounts of data, because the training set all had or all did not have the trait. No results are included for SVMs at scales where any trials failed. LightGBM, SVM, and differentiable logic require either too much time or too much memory with 70, 000 samples, and are not run on these data sets. As we can see, the majority methods eventually reach a similar score, given enough data.

**Figure 4:**
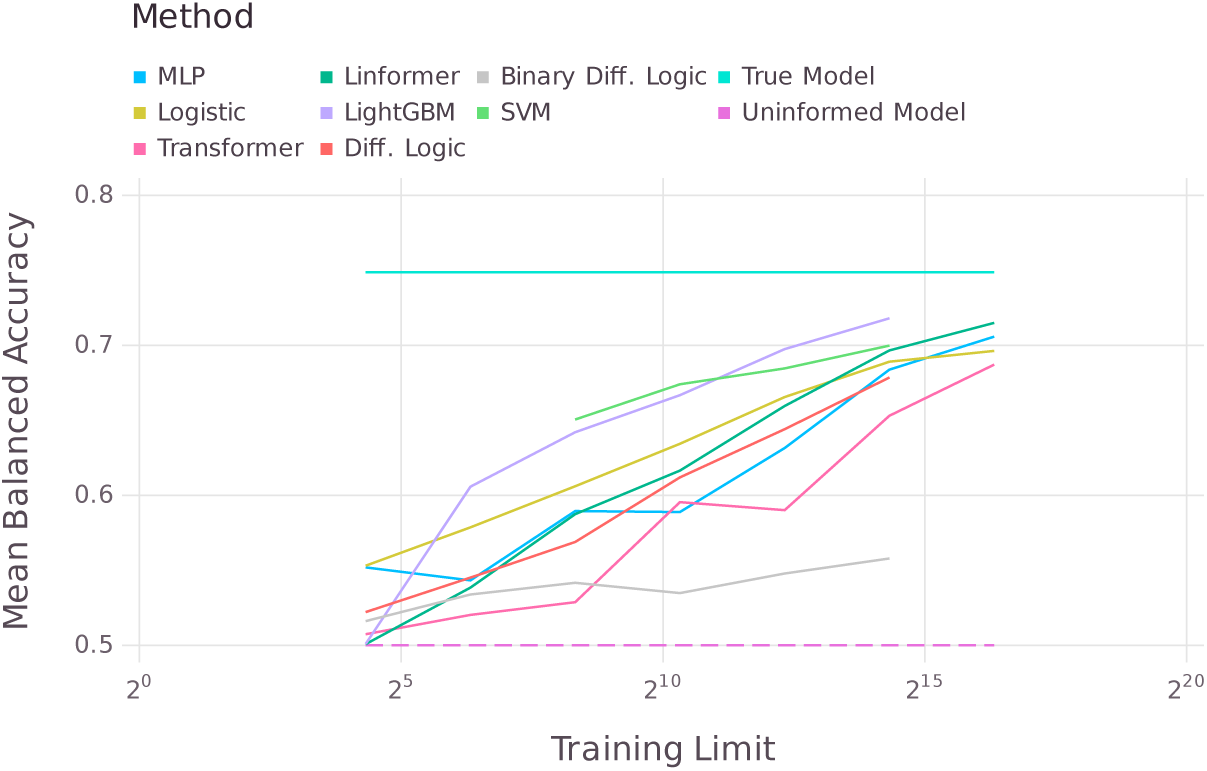
Balanced accuracy vs. training limit for each method.

With relatively little training data SVMs and LightGBM stand out, significantly outperforming all other methods. LightGBM achieves the overall best balanced accuracy, using only ≈ 20, 000 training samples. It is worth noting, however, that training on this scale required over 200GB of memory. Transformer networks with scaled-dot-product self-attention are consistently outperformed by those using Linformer, and Linformer-based networks achieved the second-best overall performance. This is likely a result of the transformer’s long running time, allowing a more thorough hyperparameter search for Linformer. While differentiable logic had generally middling accuracy, comparable to MLPs, the binarised model was only marginally better than random guessing. This suggests that the logic model is not converging to the simulated function.

### 3.2 Noise

We now investigate how the accuracy of these methods scales with increasing training size in scenarios with different levels of noise. We repeat the analysis from Section 3.1, separating the cases with no noise (Fig. 5a), low (20%, Fig. 5b) and high (40%, Fig. 5c) noise. Note that the noise determines the chance of returning a random value in the simulated gene functions and the final combined function. We average the results for each repeat with different training limits, including only the runs where all methods completed all data sets.

**Figure 5:**
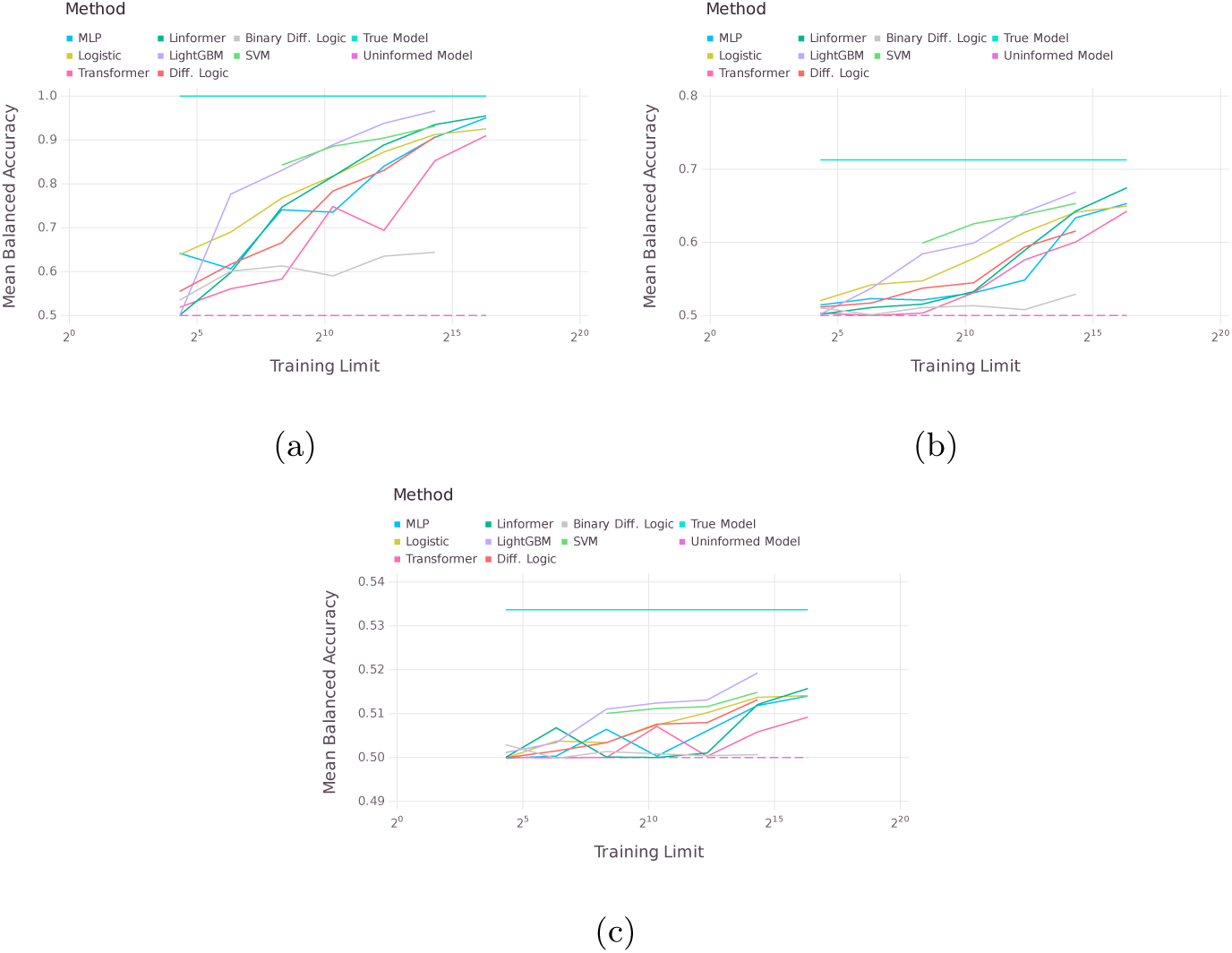
(a) Noise free balanced accuracy vs. training limit. (b) Medium (20%) noise balanced accuracy vs. training limit. (c) High (40%) noise balanced accuracy vs. training limit.

We see that without noise and with low noise, the majority of models are no better than random guessing with only 20 training samples, and approach the true model with 70, 000 training samples. While the trends are similar in both cases, it is notable that LightGBM is outperformed by Linformer when we have low noise. Increasing to 40% noise, no models are significantly better than random guessing, although the margin between the true model and random guessing is narrow here.

### 3.3 Function Complexity

To see how each method scales with increasingly complex functions, we adjust both the number of SNVs used in each gene function and the number of gene functions. Again, repeats with different training limits are averaged, including only runs where all methods completed all data sets. In this case we consider only runs without noise. We plot the mean balanced accuracy across all datasets with a given number of SNVs per gene (Fig. 6a) and a given number of regulated genes (Fig. 6b). Increasing the number of SNVs per gene decreases the performance of all methods. The only significant deviation is seen with only 2 SNVs per gene, in which case SVM achieves near perfect performance, while LightGBM is behind both SVM and Linformer. Increasing the number of simulated genes more gradually reduces performance across all methods, except for binarised differentiable logic, which performs consistently poorly.

**Figure 6:**
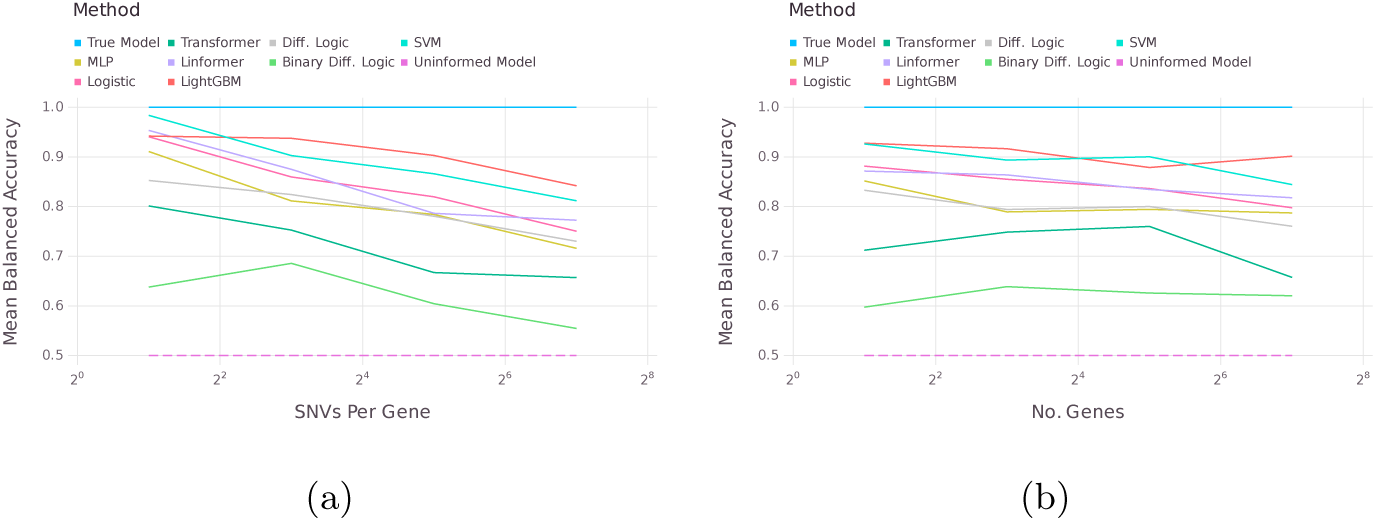
Balanced accuracy as the simulated function complexity increases. (a) Increasing regulatory SNVs per gene. (b) Increasing number of genes involved in logical function.

### 3.4 SNV Attribution

Interpretability is essential for medical applications of machine learning models [68], making it important that we are able to identify the input SNVs responsible for trait predictions. To that end we compare a number of general-purpose and transformer-specific methods for attributed predictions to particular inputs, evaluating the accuracy of each. The simplest approach, self-attention scores, do not generally explain the behaviour of the network [24]. Nonetheless, attention identifies significant patterns in some tasks [4], including in genomics [10], and may be useful in identifying significant SNVs.

To determine whether self-attention scores identify the relevant SNVs in our simulation, we take transformer networks trained on 81, 920 samples, then run one round of inference on the verification set, recording attention scores for each SNV. We add attention scores for every sample in the verification set, then calculate the Pearson correlation between the total attention score and presence in the simulation. Since the meaning of positive and negative values are unclear, we repeat the process using the absolute value of attention so that large positive and negative scores do not cancel each other.

As well as self-attention scores, we use a variety of input attribution methods implemented in captum [30]. We used Layer Integrated Gradients [65], Gradient SHAP [36], Saliency [62], Guided Backprop [63], Deconvolution [74], Input × Gradient [61], and Deeplift SHAP [36] to attribute class predictions to the encoded SNV sequence for samples in the test set. Due to their high memory requirements, we used only the first 20 samples for Layer Integrated Gradients, and the first 100 for Gradient SHAP, all other methods were run with 1, 000 samples. We then take the mean attribution score for each SNV, and calculate the Pearson correlation between that score and a binary value indicating whether the SNV had a true non-zero effect. We calculate two scores for each method, one adding the scores directly 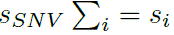, and the other adding the absolute values 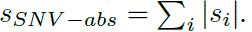. These scores were calculated separately for each simulated data set, and the mean correlation for each method is shown in Table 1.

**Table 1:** Mean correlation between attribution scores and true non-zero effect for SNV.

| Method | Mean Correlation |
| --- | --- |
| Attention | 0.013 |
| LIG | −0.020 |
| Gradient SHAP | −0.013 |
| Saliency | 0.104 |
| Guided backpropagation | 0.010 |
| Deconvolution | 0.011 |
| Input × gradient | 0.002 |
| DeepLift SHAP | 0.002 |
| Absolute attention | 0.005 |
| Absolute LIG | **0.129** |
| Absolute gradient SHAP | 0.110 |
| Absolute deconvolution | 0.105 |
| Absolute saliency | 0.104 |
| Absolute guided backpropagation | 0.105 |
| Absolute input × gradient | 0.105 |
| Absolute DeepLift SHAP | 0.105 |

Of the tested methods, none significantly deviated from zero when directly adding sores. Using absolute values instead most methods had mean correlations around 0.1. Notably, self-attention scores were not significantly correlated in either case, while layer integrated gradients with absolute values were the best-performing (mean correlation of 0.13). As we see in Fig. 7, even for the best-performing attribution method there is no dataset with a correlation above 0.4.

**Figure 7:**
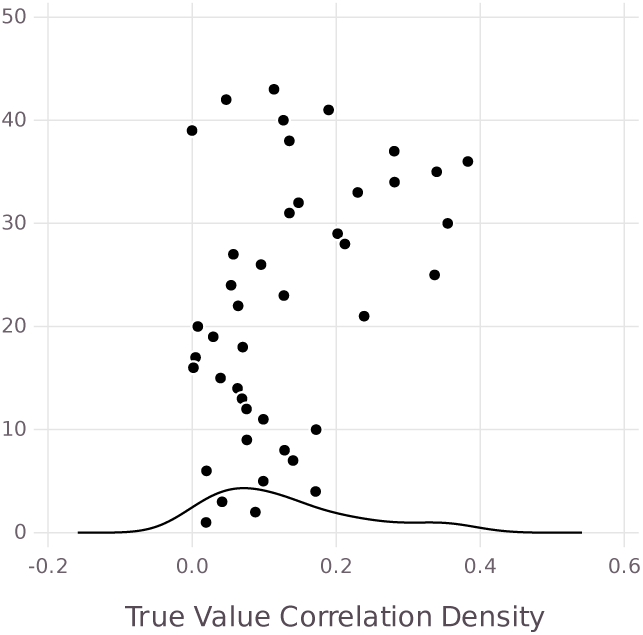
Pearson correlation between total self-attention score for each SNV and true significance of that SNV using absolute Layer integrated Gradients. Each point is the correlation in a separate simulated data set.

## 4 Alternative Transformer Architectures

As we saw in Section 3.1, Linformer-based transformer models were the most accurate method scaling to 70, 000 samples. Given the effectiveness of transformer-based models in these simulations, we explore several different Transformer variants, comparing Linformer, Informer[75], Flowformer[73], and Inflowformer, focusing on the largest training sets.

Informer reduces the time complexity to O(L log L) by adopting a sparse self-attention mechanism called ProbSparse to more effectively handle long sequence dependencies. Furthermore, Informer employs a self-attention distilling structure that effectively highlights dominant attention by halving the cascading layer inputs, enabling it to efficiently handle long input sequences. In order to linearize the quadratic time complexity of the attention mechanism in the traditional Transformer, Wu et al. proposed a flow attention mechanism with linear time complexity in the paper [73], using the conservation property of information flow from the perspective of network flow (explicitly defining the source and sink of information flow to avoid introducing competition in the attention mechanism). In addition, the flow attention mechanism does not introduce special inductive bias. When processing long sequences, time series, vision and natural language tasks, Flowformer shows strong performance, and the operation remains linear in time. Inflowformer is a hybrid model that combines the methods of Informer and Flowformer. It adopts the structural framework of Informer and replaces the ProbSparse self-attention mechanism with the Flow-Attention mechanism, aiming to explore whether this combination can demonstrate better performance in experiments compared to Informer or Flowformer alone.

We automated the hyperparameter search for each model using Ray Tune [55], covering hyperparameters including: number of layers (1 to 8), number of attention heads (1 to 16), feedforward network dimensions (128, 256, 512, 1024, 2048), learning rates from 10*^−^*^7^ to 10*^−^*^1^, and dropout rates (0.0 to 0.2). The hyperparameter search process ran for 24 hours, with the final parameter configurations presented in Table 2. We fixed all models’ batch size to 16 and embedding size to 32.

**Table 2:** Hyperparameter settings.

| Parameter | Linformer | Informer | Flowformer |
| --- | --- | --- | --- |
| Num_layers | 1 | 8 | 4 |
| Heads | 4 | 8 | 8 |
| Dim_feedforward | 256 | 1024 | 256 |
| Learning rate | $6.178 \times 10^{-5}$ | $1.164 \times 10^{-4}$ | $2.135 \times 10^{-5}$ |
| Dropout | 0.0038 | 0.0831 | 0.0224 |

**Table 3:** Construction of combined binary matrix, including age and SNVs.

| Age | Age $\geq x_1$ | Age $\geq x_2$ | ... | Age $\geq x_k$ | ... | SNV<sub>1</sub> = 0 | ... |
| --- | --- | --- | --- | --- | --- | --- | --- |
| 55 | 1 | 1 | ... | 0 | ... | 1 | ... |
| 41 | 1 | 0 | ... | 0 | ... | 0 | ... |

We use the same training, and test sets as the rest of the simulation, with hyperparameters tuned using validation set taken from the training data (containing of 7% of the overall data). We trained each model for 60 epochs, evaluating performance on the validation set after each, then using the best performing epoch’s parameters for the final evaluation on the test set.

Figs. 8 and 9 Show the balanced accuracy and time taken for each model, respectively, using 70, 000 training samples and varying the amount of noise in the data. While there are minor differences in the balanced accuracy across methods, results are generally comparable across all approaches. Linformer is notably the best-performing in two out of three tests, and the runner-up in the other. Linformer is by far the fastest attention mechanism tested, taking 10 to 25× less time than Informer, Flowformer, or Inflowformer.

**Figure 8:**
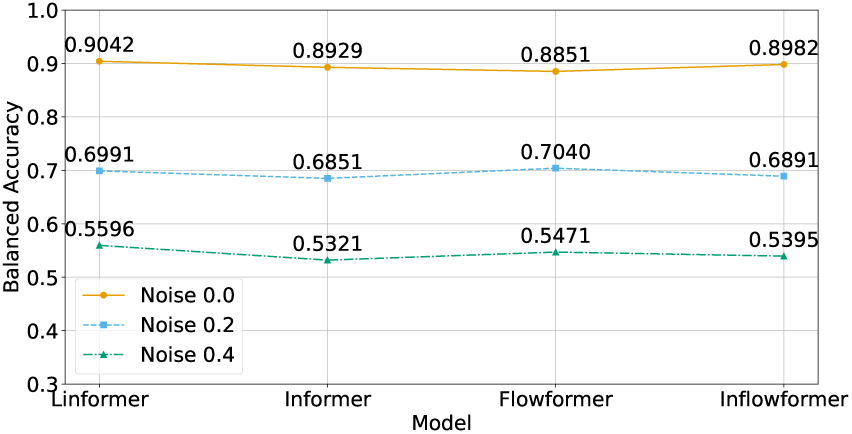
Balanced accuracy of transformer-based models at varying noise levels.

**Figure 9:**
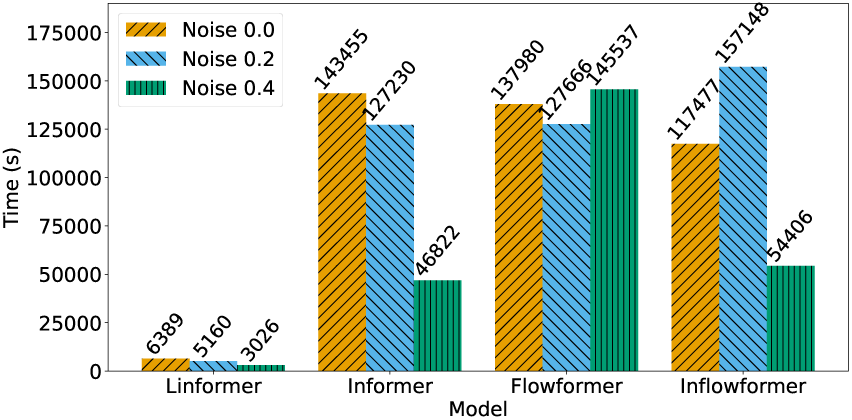
Time taken for Transformer-based models to achieve their best balanced accuracy.

Given its significant performance advantage and comparable accuracy, we will focus on Linformer-based attention models from here on.

## 5 Gout Prediction in the UK Biobank

GWAS have identified common SNVs contributing to gout. These do not fully explain the heritability of gout, however [38, 37]. Building on this work, we test several of the most scalable methods from Section 3 ability to identify nonadditive effects that might explain some of the missing heritability [39].

### 5.1 Data

Here we use UK Biobank (UKBB) (project 12611) data to predict gout and identify predictive genetic factors. UKBB is one of the largest human genome data sets to date [64]. The genetic data includes a set of ≈ 850, 000 SNVs collected using the UK Biobank Axiom Array, across ≈ 488, 000 patients. There is an additional set of ≈ 90 million variants imputed using the Haplotype Reference Consortium and UK10K and 1000 Genomes reference panels. These imputed variants are not measured directly, but are strongly correlated with SNVs that were included in the genotyping array and can be inferred with reasonable confidence (over 90% accuracy for SNVs with non-reference alleles occurring in 2% of the population [69]). The UKBB also contains measurements of serum urate levels (µmol/L), as well as gout diagnoses, age, sex, and BMI for each patient. Age, sex, and BMI are all known to be highly predictive of gout [66, 32], and sustained high levels of serum urate are necessary (although not sufficient) for developing gout [12]. As well as developing an accurate gout classifier, we aim to identify the predictive genetic factors with these models. We include age, sex, and BMI in our data to account for any interaction between these and genetic risk factors, as well as to avoid conflating correlated genomic variants.

We reduce the data to a subset of 13, 290 SNVs by taking only variants associated with urate with *p* ≤ 0.05 according to a recent GWAS study [37]. The resulting filtered set contains ≈ 488, 000 individuals, of which only ≈ 9, 000 have gout. To avoid training with an unbalanced data set, we use only a random subset of the non-gout cases in each training epoch, matching the number of gout cases used. Across all methods, we use only the first 70% of samples for training. We use 15% as a validation set for hyperparameter selection and reserve 15% as a test set.

Several other reductions of the dataset were also considered, these are described in Appendix B.

### 5.2 Methods

We tested logistic regression, Pint, differentiable logic, and transformers for predicting gout using 13, 290 SNVs from the UK Biobank, as well as age, sex, and BMI. We were not able to run either gradient-boosted decision trees or support vector machines on this dataset. The two best-performing approaches, lasso regression and a transformer encoder, are described briefly here. Hyperparameter details for each method can be found in Appendices C to F.

#### 5.2.1 Pairwise Lasso Regression

To account for possible epistasis, we use the Pint lasso regression method [17] that includes a term for every pairwise interaction between variables. We construct a binary matrix for each demographic *ρ_i_* by choosing *k* thresholds *v_i,_*_1_*, …, v_i,k_*, then using *k* binary variables in the place of *ρ_i_* to encode whether *ρ_i_* was at least *v_i,k_*. Specifically, 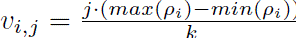. We then replace each sample *ρ_i_* with the binary variables *b_i,_*_1_*, …, b_i,k_*, where *b_i,j_* = 1 if and only if *ρ_i_* ≥ *v_i,j_*. In the following analysis we use *k* = 10 for both age and BMI. Sex is 1 if the sample is recorded as male, 0 otherwise. SNVs are encoded using encoding V5 (see Appendix F.1 for details), giving us an SNV matrix *X* ∈ {0, 1, 2}*^n×p^*. We convert this ternary matrix to binary by replacing each column *X_j_* with three columns 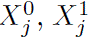 and 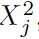, with an entry 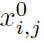 of 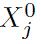 equal to one if and only if *x_j_* = 0. Similarly, 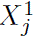 and 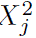 encode whether the original column was 1 or 2. Finally, we construct a combined genotype-demographic matrix by concatenating columns of the binary SNV and demographic matrices. Table 5 shows an example of a combined matrix using only age and SNVs.

Since Pint is a linear regression model, we cannot directly use it for classification. Nonetheless, we can predict serum urate, and then use the predicted urate level as a predictor of gout. While the test and validation sets are the same as other methods, with Pint we use an unbalanced training set containing 377, 605 samples. We tested with and without interactions, with 500 or 10, 000 non-zero effects allowed, with and without the approximate hierarchy assumption, with and without age, sex, & BMI, and predicting both urate and *log*(urate). The best-performing model in the validation set used pairwise interactions, 10, 000 non-zero effects, included age, sex, & BMI, assumed a hierarchy was present and fit urate directly. For further details on the Hyperparameter selection process see Appendix D.

#### 5.2.2 Transformer Encoder

We use a variation of the SNVformer architecture [18], with additions to the SNV encoding. We tested several variations of the original SNVformer encoding (see Appendix F.2) the best-performing variation is described here. Each SNV is encoded in four components, with separate embeddings for the nucleotide change, chromosome, position, and gene (Fig. 10).

**Figure 10:**
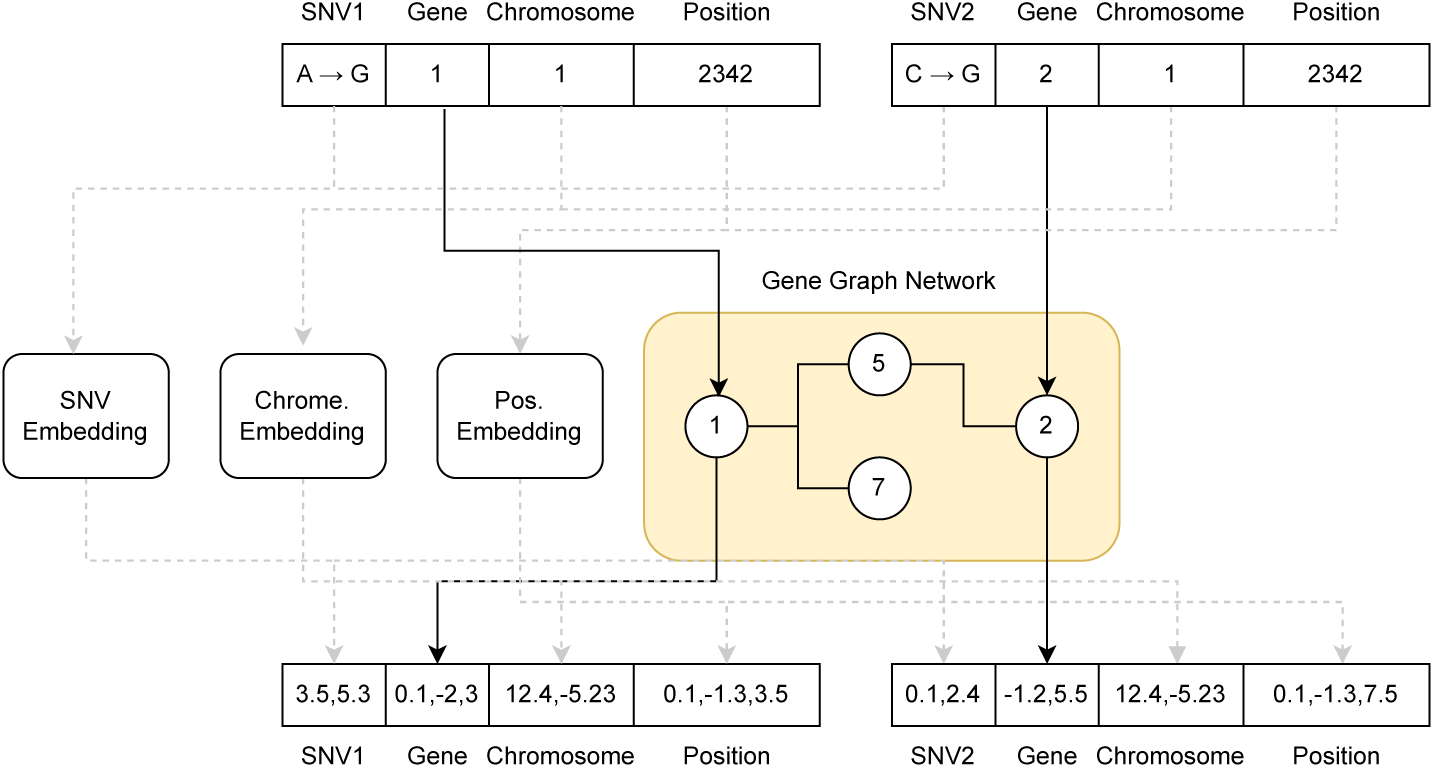
SNV embedding with graph neural network gene encoding.

**Gene and Chromosome Embedding** For every SNV, we optionally include a learned embedding of its gene and chromosome, if applicable. This allows us to identify relationships between genes, and associate SNVs in the same gene or chromosome. Particularly in pre-training, this should lead to higher attention between SNVs in correlated genes. We do not need to learn these relationships from scratch, however.

We either use a simple learned embedding, or optionally encode genes using a Graph Convolutional Network [29, 20], with a node for each gene present in the dataset. Edges are placed between nodes when an interaction of any kind between the corresponding genes is present in the BioGRID interaction database [46]. Given the graph adjacency matrix *A*, input gene embedding *X*, and learnable weight parameters *W*_0_ and *W*_1_, the graph network outputs a vector *Z* ∈ R*^n×d^* where the embedding of every gene is influenced by all genes connected in the graph.

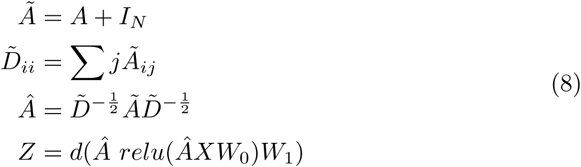

We use the same embedding dimension *d_embed_*___*_genes_* for nodes in the graph network and hidden layers. These output embeddings retain the input dimensions *n_genes_* × *d_embed_*___*_genes_*, and are used instead of the randomly-initialised learned embeddings of the simple gene encoding above. The complete SNV encoding with gene-graph encoding is shown in Fig. 10.

Using this SNV embedding, we train an encoder-only transformer (Fig. 11) to classify gout cases. Our transformer is implemented using Linformer selfattention [72] with the memory efficient attention implemented from PyTorch 2.0 [49]. We use a hyperparameter search to choose the exact architecture of the transformer.

**Figure 11:**
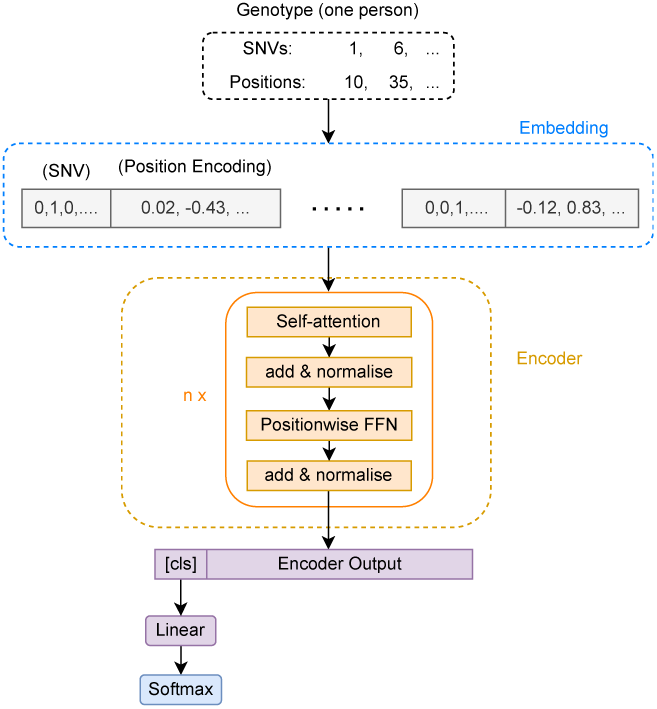
Encoder-only transformer architecture

### 5.3 UKBB Results

We see a general trend with all models, where the best hyperparameter choices result in test-set cross-entropy loss around 0.5, and AUROC around 0.8. Nonetheless, performance of a tuned transformer network was similar to the best-performing linear model (0.832 for transformers vs. 0.836 for regression). Both options significantly outperformed differentiable logic. It is worth noting that Pint achieved very similar performance both with and without pairwise interactions, AUROC of 0.836 and 0.837 respectively. This suggests that it is the choice of lasso-regularised regression, rather than the inclusion of interactions, that is beneficial here.

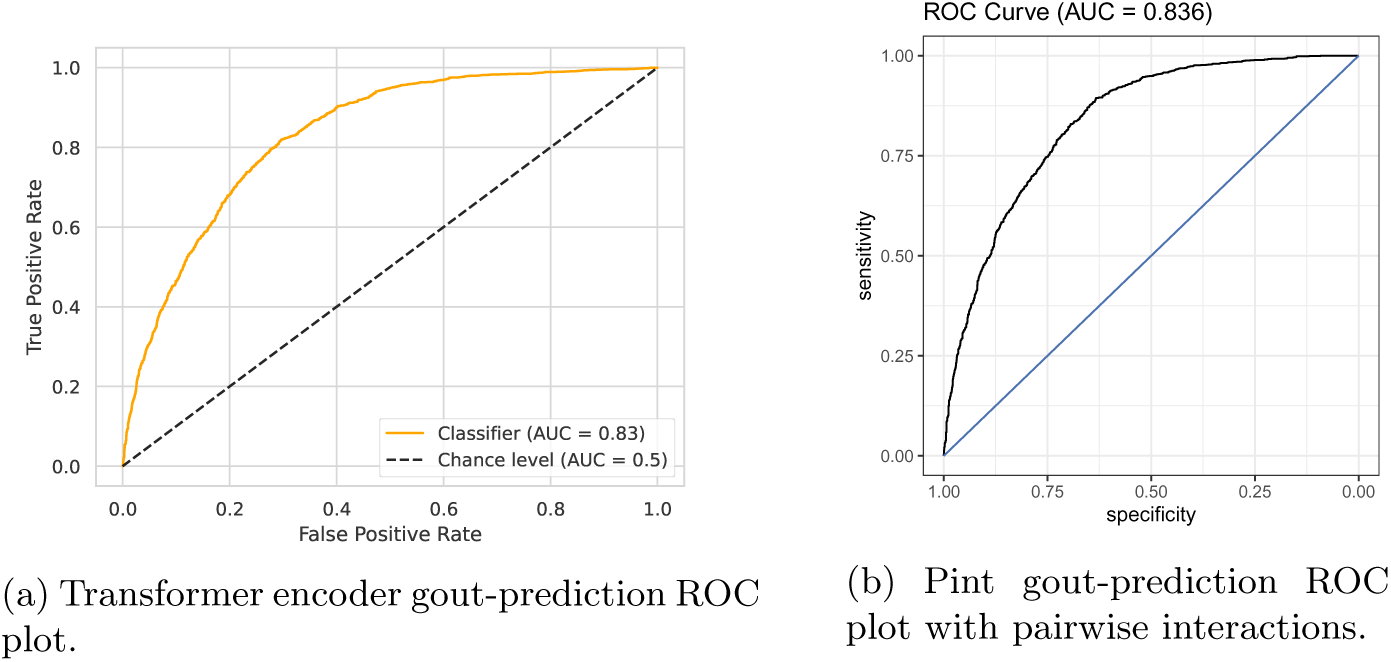

## 6 Discussion

We saw in Section 3 that most methods converge to similar balanced accuracy as the amount of training data increases, a trend that is similar regardless of noise (Section 3.2). Nonetheless, there are significant differences in both the computational resources, and training data required between methods. Logistic regression, LightGBM, and SVMs, have an advantage with only a few hundred training samples and no noise. While logistic regression remains computationally tractable at large scales, LightGBM and SVMs require too much time or memory. Even limited to ≈ 20, 000 training samples, LightGBM is the best performing method overall, however. Linformer-based transformers require over 16, 000 samples to outperform regression, but were the best-performing method that scaled to 70, 000 training samples. Binarised logic models performed consistently poorly, and classic Transformers were consistently outperformed by Linformer, although this may be due to Linfromer trying more hyperparameters within the time limit. Interestingly, with enough data linear regression and SVMs perform nearly as well as non-linear methods, despite the simulated functions not necessarily being linearly separable. While transformer-encoders performed well as predictors, it is worth noting that none of the SNV attribution methods we tested produced scores that were highly correlated with the ground truth (Section 3.4).

Notably, while there is a set of parameters for the differentiable logic model that perfectly matches the simulated function, overall performance is not exceptional, remaining broadly comparable to that of MLPs. Given the difference between fuzzy logic and binarised model performance (Fig. 5), it is likely that the fuzzy model is not trending towards a binary logical function, but rather finding a locally optimal approximation that relies on the functions ∧(*a, w*), ∨(*a, w*) having non-binary weights. Since these do not correspond to functions in the true simulated function, except when their weights are binary, this approximation has no particular advantage over the other methods.

We applied a number of models, including regression and transformer encoders, to predicting gout in the UK Biobank. Both lasso regression and transformer-encoders performed similarly as gout predictors, with results comparable to the current state-of-the-art. Despite this, neither regression with interactions nor the more powerful transformer encoders outperformed classic approaches. While there was a marginal improvement logistic regression (Table 4), regularised regression (Pint) remains the best-performing approach. Lasso-regularised regression without interactions remained the best-performing method. This suggests that we either do not have enough samples to fit a complex model, or a significant number of the genetic causes were not included in the UKBB genotype panel. The first is particularly likely, since we only have around 9, 000 individuals with gout in our data, and both models perform comparably at this scale in our simulations. While including more positive training samples would be the ideal solution, we could instead attempt to overcome this limitation with data augmentation or inclusion of prior information. While we attempted a simple data-augmentation approach in Appendix F.10 without success, another approach may yield better results. Some prior information was also included in the transformer gene embedding graph, however this does not appear to have significantly influenced the network. An alternative approach would be to use prior information to regularise the network, directly encouraging attention between genes with known associations and penalising others in the loss function. A similar approach could be used to encourage attention to known gout or urate associated genes and SNVs, penalising but still allowing attention elsewhere. While this would restrict the model’s ability to detect entirely unknown interactions, such regularisation could reduce overfitting and improve generalisation. It is worth noting that the best performing method was lasso regression (Table 4), which is also a heavily regularised model.

**Table 4:** Best validation set AUROC for each method.

| Method | Best AUROC |
| --- | --- |
| Logistic | 0.825 |
| Pint | **0.836** |
| Differentiable Logic | 0.790 |
| Transformers | 0.832 |

There are a number of opportunities to expand on this work. While our analysis of Gout in the UK Biobank did consider age, sex, and BMI, there are other non-genetic factors known to influence gout risk [58]. By more including all relevant non-genetic factors it is likely that a future study could produce a more accurate model of gout risk. Similarly, we restricted our input to a relatively small subset of SNVs to reduce the computational cost of training. While these were chosen for their association with either gout or urate, with sufficient computational resources a model could be trained on the entire unimputed dataset or even the whole genome, which would allow accounting for the effects of rare variants. The particular combination of Linformer and memory efficient attention used may not be the most efficient approach. There are many alternatives to the transformer architecture for long sequence data, one of which may be more effective. While we considered transformers both with and without pre-training (Appendix F.8), there was no significant difference in the gout-prediction performance between the two. More computationally efficient models would also allow significantly longer pre-training, which may have a larger effect. Finally, while our approach in Section 5.2.2 used known gene-gene interactions in the gene embedding, this had only a small impact on validation-set accuracy. Alternative approaches to including prior information in the model may be more effective and are worth exploring.

In our simulation study (Section 3) transformer networks had a marginal advantage over regression in predicting simple non-linear traits. Despite the functions including non-linear logical components, regression was able to find close approximations in the majority of cases. In more complex simulations where regression is not able to closely approximate the function there might be a more significant difference between methods, and designing and running such a simulation would be worthwhile.

## 7 Acknowledgementes

This work was partially supported by Royal Society Te Apārangi through a Rutherford Discovery Fellowship [UOC1702] and a Marsden grant [21-UOC057]; and Ministry of Business, Innovation, and Employment of New Zealand through an Endeavour Smart Ideas grant [UOOX1912] and a Data Science Programmes grant [UOAX1932].

This research was funded in part through a Fred Hutchinson Cancer Center Translational Data Science Integrated Research Center Postdoctoral Fellowship.

This work uses data provided by patients and collected by the NHS as part of their care and support.

We gratefully acknowledge all data contributors, i.e., the Authors and their Originating laboratories responsible for obtaining the specimens, and their Submitting laboratories for generating the genetic sequence and metadata and sharing via the GISAID Initiative, on which this research is based. This research has been conducted using the UK Biobank Resource under application number 12611.

