## Appendix for "Accuracy and Scalability of Machine Learning Methods for Genotype-Phenotype Association Data"

### A Simulation Parameters

We simulate 48 datasets using the method described in Section 1, with varying levels of noise and degrees of complexity. We use 2, 8, 32, and 128 SNVs in each expression regulatory function (Equation 2). For each of these, we simulate sets with 2, 8, 32, and 128 genes in the logical trait function (Fig 1). For each combination of SNVs per gene and genes per trait, we simulate a set with no noise, one with 20%, and one with 40% noise.

#### A.1 Hyperparameter Optimisation

We run a separate hyperparameter search for each simulated dataset. We allow the number of SNVs included in each linear function to be between 1 and 32, the number of linear functions between 1 and 32, the number of epochs between 1 and 32, and choose learning rates between $10^{-1}$ and $10^{-7}$ on a log scale.

**Logistic Regression** We run a hyperparameter search using Ray Tune [55] with the AHSA scheduler [31] to decide the learning rate and dropout hyperparameters. Learning rate is allowed to vary between $10^{-7}$ and $10^{-1}$ on a log scale, and dropout between 0.0 and 0.2. After each epoch we calculate the error on the validation set, and finally the model with the best validation error is used for evaluation. While regression cannot model the non-linear logical component of the simulation, we use this method as a baseline for comparison.

**Gradient-Boosed Decision Trees** Hyperparameters are chosen using a grid search with a target of 100 trials, implemented in MLJ [5], allowing the following hyperparameter ranges: the minimum number of samples identified by each leaf ranges from 10 to $10,000$, sampled evenly on a log scale; the maximum tree depth is evenly sampled from 2 to 20; learning rate ranges from $10^{-4}$ to 1 on a log scale; the number of leaves ranges from 1 to 1024. Among all trials the best is chosen using 3-fold cross-validation on the training set.

**Multi-Layer Perceptron** The number of hidden neurons per layer and depth are chosen using Ray Tune. Hidden neurons vary from 2 to 256, and depth from 1 to 16, both sampled on a log scale.

**Transformer Encoder** Both Linformer and Transformer architectures are optimised using Ray Tune. The following hyperparameter ranges are used for the Linformer. The learning rate varies between $10^{-7}$ and $10^{-1}$ on a log scale, and dropout is between 0.0 and 0.2. We use between 1 and 8 encoder layers, each with 1, 2, 4, or 8 attention heads. The SNV embedding size is 16, 32, 64, 128, or 256. The Linformer k value is 16, 32, 46, 128, or 256. The feedforward hidden layer has a size of 128, 256, 512, 1024, 2048, or 4096.

For the transformer, we use one or two encoder layers, with one or two attention heads. The SNV embedding size is 8, 16, 32, 64, or 128. Finally, the feedforward hidden layer has size 128, 256, 512, 1024, or 2048. Note that transformer self-attention requires significantly more time and memory than Linformer self-attention, and as a result the network has to be significantly smaller in other respects. The structure of the transformer does not suggest any particular way in which it might fit the simulation. Nonetheless, both linear and logical functions are able to be represented by the internal components, in the same manner as MLPs.

**Differentiable Logic** Hyperparameters for the differentiable logic model are chosen using Latin Hypercube sampling [2] and the best-performing model on a 10% hold-out validation set is then used in the test set. We also test the binarised version of each fitted model, using the scheme described in Appendix E.3. With the correct parameters this model can perfectly implement the simulated function. How successful the approach is in simulations will therefore depend primarily on the ability to choose these parameters with random hyper-parameter searches and the AdamW algorithm.

##### A.1.1 Hardware and Time Constraints

The differentiable logic model, and support vector machines are all fit using four threads each on a Intel(R) Xeon(R) Gold 6342 CPU, with $\approx$ 500GB of memory shared between four simultaneous hyperparameter searches. Regression and MLP models are run on the same system using one NVIDIA RTX A6000. Transformers, including Linformer, are fit using 32 threads on an AMD EPYC 7713 and four NVIDA A100 GPUs. The regression and MLP hyper-parameter searches are allowed to run for up to 600 seconds, except when 81920 training samples are used, in which case 1200 seconds are allowed. The Linformer hyper-parameter search is allowed $\frac{600+n_{train}}{16}$ seconds, where $n_{train}$ is the number of training samples, and the classic transformer is allowed $\frac{1200+n_{train}}{4}$ seconds. SVM, and differentiable logic hyperparameter searches have no time limit.

### B Reducing UKBB Dataset Size

The entire UK Biobank contains too many SNVs to use directly with any of our non-linear methods. We instead construct smaller datasets in a number of ways, either by targeting only the variants most likely to be relevant to gout (Appendix B.1), or by avoiding including redundant information (Appendix B.2).

#### B.1 GWAS-Based Filtering

To begin with, we re-use the GWAS results from [37] to identify the association between each SNV and serum urate using a genome wide association study (GWAS).

Given this GWAS, we then take only SNVs associated with serum urate with a p-value of at most 0.1 to form the set *GWLarge* of 65,803 variants. We also produce a smaller set of only $13,290$ SNVs with p-values $\leq 0.05$ called *GWMed*. Finally, we have a set *GWSmall* with only 123 SNVs identified in another GWAS by Tin et al. [66].

not used in main paper

##### B.2 Linkage Disequilibrium Filtering

As an alternative to relying on important SNVs being directly associated with urate (and therefore detectable with a GWAS), we reduce the data size by removing redundant information. Since the imputed variants can, by definition, be identified given the unimputed variants, we begin with only the unimputed $850,000$ SNVs. We then filtered by removing any SNVs with a pairwise coefficient of determination ($r^2$) greater than $10^{-5}$, using plink to greedily remove SNVs until no such pairs remain. The resulting set *LDMed* contains $13,228$ SNVs. Finally, we construct a set *LDSmall* by filtering out all SNVs with an $r^2 > 0.1$ from the *GWMed* set, resulting in 320 uncorrelated SNVs, each of which is associated with serum urate with a p-value of less than 0.05.

**Balanced Sets** All filtered sets contain $\approx 488,000$ individuals, of which only $\approx 9,000$ have gout. For many of our models, this unbalanced training set is a problem, as it results in a strong bias towards non-gout predictions. Unless otherwise stated, we avoid this by training and testing using balanced subsets of the data, containing all gout cases and an equal number of non-gout cases.

#### C Elastic Net Regularised Regression

We use Linear regression with elastic net regularisation, implemented in the Julia package MLJLinearModels [40] as a baseline. We use the 123 SNVs present in the UK Biobank that were identified by Tin et al. [66] as being associated with urate, along with age, sex, and BMI.

We construct a binary matrix for each demographic $\rho_i$ by choosing $k$ thresholds $v_{i,1}, \ldots, v_{i,k}$, then using $k$ binary variables in the place of $\rho_i$ to encode whether $\rho_i$ was at least $v_{i,k}$. Specifically, $v_{i,j} = \frac{j \cdot (max(\rho_i) - min(\rho_i))}{k}$. We then replace each sample $\rho_i$ with the binary variables $b_{i,1}, \ldots, b_{i,k}$, where $b_{i,j} = 1$ if and only if $\rho_i \geq v_{i,j}$. In the following analysis we use $k = 10$ for both age and BMI. Sex is 1 if the sample is recorded as male, 0 otherwise. SNVs are encoded using encoding V5 (see Appendix F.1 for details), giving us an SNV matrix $X \in \{0, 1, 2\}^{n \times p}$. We convert this ternary matrix to binary by replacing each column $X_j$ with three columns $X_j^0$, $X_j^1$ and $X_j^2$, with an entry $x_{i,j}^0$ of $X_j^0$ equal to one if and only if $x_j = 0$. Similarly, $X_j^1$ and $X_j^2$ encode whether the original column was 1 or 2. Finally, we construct a combined genotype-demographic matrix by concatenating columns of the binary SNV and demographic matrices. Table 5 shows an example of a combined matrix using only age and SNVs.

| Age | Age $\geq x_1$ | Age $\geq x_2$ | ... | Age $\geq x_k$ | ... | $SNV_1 = 0$ | ... |
|---|---|---|---|---|---|---|---|
| 55 | 1 | 1 | ... | 0 | ... | 1 | ... |
| 41 | 1 | 0 | ... | 0 | ... | 0 | ... |

Table 5: Construction of combined binary matrix, including age and SNVs.

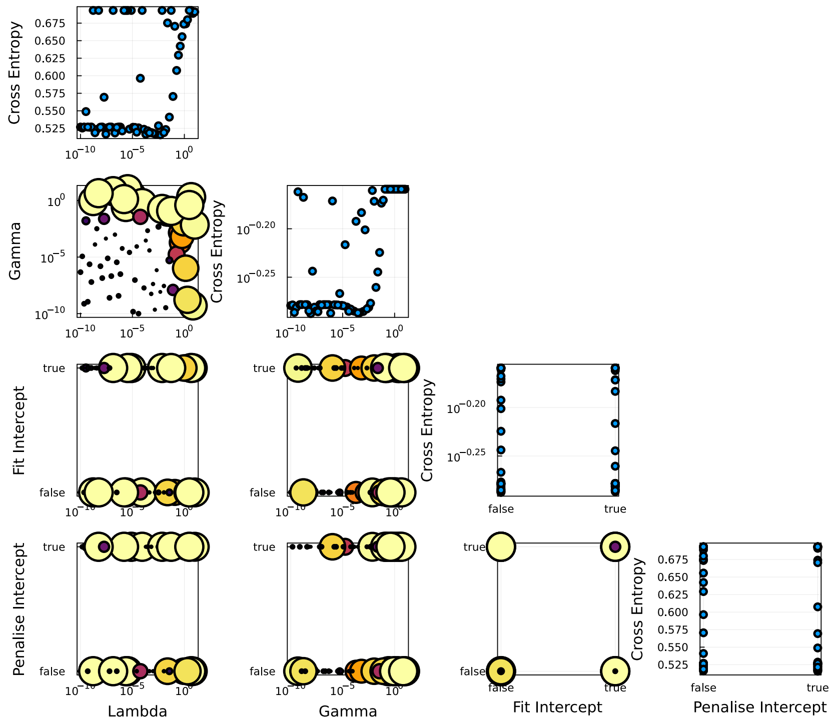


Figure 13: Elastic net hyperparameter search. Cross entropy for each trial is shown, separated by hyperparameter. Parameter vs. parameter plots show the distribution of hyperparameter samples. Point size and color in parameter vs. parameter plots indicate test-set loss of the trial, with smaller darker points being more successful (lower loss).

Hyper-parameters are chosen using the Latin Hypercube [2] hyperparameter search implemented in MLJ [5]. The results of our hyperparameter search are shown in Fig. 13. Low cross entropy loss is only achieved with lambda and gamma both below 1, and we see significant improvements when either is below $10^{-2}$. With low enough values most models perform similarly, however. There is no clear advantage to including or penalising the intercept (Fig. 13, cross entropy vs. penalised intercept). The best-fitting model achieves an AUROC of 0.825 in the validation set (Fig. 14), and accuracy of 0.742. This result is comparable to the state-of-the-art (AUROC 0.84) by Tin et al. [66].

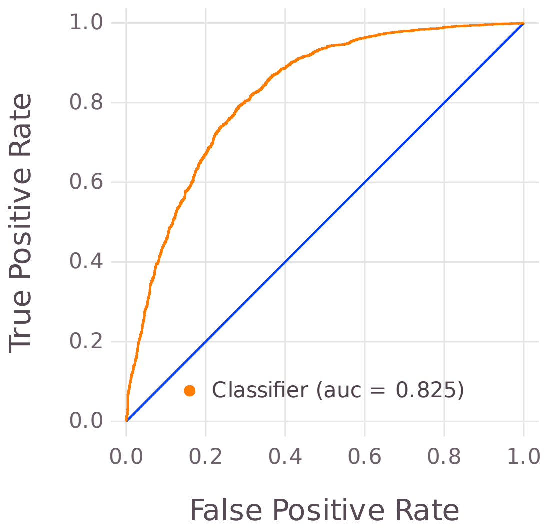


Figure 14: Elastic net regression ROC curve

### D Modelling Gout with Pint

Since Pint [17] is a linear regression model, we cannot directly use it for classification. Nonetheless, we can predict serum urate, and then use the predicted urate level as a predictor of gout. We test each of the datasets described in Appendix B.1, in each case using 70% of gout samples for training, and the remaining 30% for testing and validation, with an equal number of non-gout samples in the testing and validation sets. Note that unlike other methods in this section, Pint does not use only a balanced subset of the training data. While the test and validation sets are the same as other methods, Pint uses an unbalanced training set containing $377,605$ samples.

**Model Design** Using a binary matrix constructed with the method in Appendix C, we separately predict both urate (µmol/L) and $log$(urate). Using both urate and log urate allows us to model additive and multiplicative SNV effects, respectively. Urate (or log(urate)) predictions are then used as predictors of gout. We construct the ROC Curve and find the area under the curve for each choice of parameters. Results for all trials are shown in Table 6, with hyperparameters shown on the left, training, test and validation set mean squared error for urate and validation set gout prediction AUROC are shown on the right.

| Dataset | Depth | Max. $\neq 0$ | Hier. | Geno. | Demo. | $log(urate)$ | Val. Loss | Train loss | Test Loss | Test AUROC |
|---|---|---|---|---|---|---|---|---|---|---|
| *GWMed* | 1 | 10000 | true | true | false | false | 3.155 | 1.631 | 3.343 | 0.668 |
| *GWMed* | 2 | 10000 | true | true | false | false | 3.155 | 1.613 | 3.357 | 0.660 |
| *GWMed* | 1 | 10000 | true | true | false | true | 0.093 | 0.062 | 0.098 | 0.667 |
| *GWMed* | 2 | 10000 | true | true | false | true | 0.094 | 0.062 | 0.098 | 0.651 |
| *GWMed* | 1 | 10000 | true | true | true | false | **2.147** | 0.978 | 2.239 | **0.837** |
| *GWMed* | 2 | 10000 | true | true | true | false | **2.135** | 0.956 | 2.205 | **0.836** |
| *GWMed* | 1 | 10000 | true | true | true | true | **0.063** | 0.037 | 0.064 | 0.837 |
| *GWMed* | 2 | 10000 | true | true | true | true | **0.061** | 0.035 | 0.062 | 0.835 |
| *GWMed* | 1 | 500 | false | true | false | false | 3.248 | 1.687 | 3.405 | 0.652 |
| *GWMed* | 2 | 500 | false | true | false | false | 3.266 | 1.701 | 3.440 | 0.646 |
| *GWMed* | 1 | 500 | false | true | true | false | 2.190 | 1.024 | 2.266 | 0.830 |
| *GWMed* | 2 | 500 | false | true | true | false | 2.202 | 1.046 | 2.297 | 0.822 |
| *LDSmall* | 1 | 1000 | false | false | true | false | 2.347 | 1.143 | 2.404 | 0.808 |
| *LDSmall* | 2 | 1000 | false | false | true | false | 2.334 | 1.132 | 2.384 | 0.806 |
| *LDSmall* | 1 | 1000 | false | false | true | true | 0.068 | 0.044 | 0.069 | 0.808 |
| *LDSmall* | 2 | 1000 | false | false | true | true | 0.067 | 0.043 | 0.068 | 0.804 |
| *LDSmall* | 1 | 1000 | true | true | false | false | 3.266 | 1.699 | 3.491 | 0.623 |
| *LDSmall* | 2 | 1000 | true | true | false | false | 3.278 | 1.705 | 3.501 | 0.620 |
| *LDSmall* | 1 | 1000 | true | true | false | true | 0.097 | 0.065 | 0.103 | 0.619 |
| *LDSmall* | 2 | 1000 | true | true | false | true | 0.097 | 0.066 | 0.103 | 0.619 |
| *LDSmall* | 1 | 1000 | true | true | true | false | 2.201 | 1.037 | 2.298 | 0.826 |
| *LDSmall* | 2 | 1000 | true | true | true | false | 2.194 | 1.033 | 2.288 | 0.824 |
| *LDSmall* | 1 | 1000 | true | true | true | true | 0.064 | 0.039 | 0.066 | 0.825 |
| *LDSmall* | 2 | 1000 | true | true | true | true | 0.062 | 0.039 | 0.064 | 0.823 |
| *LDSmall* | 1 | 100 | true | true | true | false | 2.244 | 1.072 | 2.327 | 0.820 |
| *LDSmall* | 2 | 100 | true | true | true | false | 2.267 | 1.079 | 2.328 | 0.809 |
| *LDSmall* | 1 | 100 | true | true | true | true | 0.065 | 0.041 | 0.066 | 0.819 |
| *LDSmall* | 2 | 100 | true | true | true | true | 0.064 | 0.041 | 0.065 | 0.809 |
| *LDSmall* | 1 | 500 | false | true | false | false | 3.270 | 1.704 | 3.495 | 0.618 |
| *LDSmall* | 2 | 500 | false | true | false | false | 3.312 | 1.720 | 3.531 | 0.603 |
| *LDSmall* | 1 | 500 | false | true | false | true | 0.097 | 0.065 | 0.103 | 0.615 |
| *LDSmall* | 2 | 500 | false | true | false | true | 0.099 | 0.066 | 0.104 | 0.599 |
| *LDSmall* | 1 | 500 | false | true | true | false | 2.202 | 1.041 | 2.297 | 0.825 |
| *LDSmall* | 2 | 500 | false | true | true | false | 2.211 | 1.051 | 2.298 | 0.820 |
| *LDSmall* | 1 | 500 | false | true | true | true | 0.064 | 0.039 | 0.066 | 0.824 |
| *LDSmall* | 2 | 500 | false | true | true | true | 0.062 | 0.039 | 0.064 | 0.821 |
| *LDMed* | 1 | 10000 | true | true | false | false | 3.531 | 1.783 | 3.712 | 0.509 |
| *LDMed* | 2 | 10000 | true | true | false | false | 3.611 | 1.633 | 3.835 | 0.499 |
| *LDMed* | 1 | 10000 | true | true | false | true | 0.106 | 0.069 | 0.111 | 0.509 |
| *LDMed* | 2 | 10000 | true | true | false | true | 0.110 | 0.064 | 0.115 | 0.502 |
| *LDMed* | 1 | 10000 | true | true | true | false | 2.359 | 1.129 | 2.405 | 0.803 |
| *LDMed* | 2 | 10000 | true | true | true | false | 2.425 | 1.023 | 2.452 | 0.785 |
| *LDMed* | 1 | 10000 | true | true | true | true | 0.068 | 0.043 | 0.069 | 0.804 |
| *LDMed* | 2 | 10000 | true | true | true | true | 0.070 | 0.040 | 0.071 | 0.781 |
| *LDMed* | 1 | 500 | false | true | true | false | 2.352 | 1.135 | 2.399 | 0.806 |
| *LDMed* | 2 | 500 | false | true | true | false | 2.352 | 1.158 | 2.409 | 0.798 |
| *LDMed* | 1 | 500 | false | true | true | true | 0.068 | 0.044 | 0.069 | 0.807 |
| *LDMed* | 2 | 500 | false | true | true | true | 0.068 | 0.044 | 0.069 | 0.799 |

Table 6: Pint urate results. Best validation set AUROC, test set urate, and log(urate) loss are in bold. *Depth* is the maximum number of variables in an interaction term. *Max* $\neq 0$ is the maximum number of parameters $\beta_i \neq 0$. *Hier.* indicates use of the approximate hierarchy assumption from [17]. *Geno.* indicates whether SNVs were included in the model. *Demo.* indicates whether age sex and BMI were included. *Log(urate)* indicates whether $log(urate)$ was the target, as opposed to simply *urate*.

**LDSmall** Using only the *LDSmall* set of 320 uncorrelated SNVs (see Appendix B.2) with no interactions, we manage an AUROC of 0.826 predicting gout, comparable to Tin et al. [66]'s state-of-the-art AUROC of 0.84. Note that the best results targeted *urate*, not $log(urate)$, included both genotypes and demographics (i.e. age, sex, and BMI), and excluded interactions. The regularisation parameter $\lambda$ was gradually reduced until $1,000$ non-zero effects were found, then fit without regularisation using only these effects.

Allowing pairwise interactions, otherwise using the same parameters, we see a marginal reduction in test set urate loss (2.201 to 2.194), indicating better serum urate predictions. Although the difference is marginal, this suggests that epistasis may be useful for predicting hyperuricaemia. Gout classification is marginally worse, however, with AUROC of only 0.824. Note that these

SNVs were not chosen for their relevance to gout, only for their low linkage disequilibrium scores (see Appendix B.2).

**LDMed** Using low linkage disequilibrium SNVs with $r^2 < 0.00001$, we again see the best validation set results without interactions, with AUROC of 0.807. The test set urate error is again marginally lower with pairwise effects (0.0680 vs. 0.0679), and gout AUROC is marginally worse at 0.799. This again suggests epistasis may be relevant to urate, but not gout.

**GWMed** The best results are achieved fitting *GWMed*, a set that has been filtered down to only significant SNVs in the GWAS ($p < 0.1$). We gradually reduce $\lambda$ until $10,000$ non-zero variants are found, then fit without regularisation using only these variants. The fast approximate hierarchy method from [17] was used, and demographics were included. Results were similar when predicting both $urate$ and $log(urate)$, and we focus on the $urate$ results below.

Allowing only individual effects, we manage a mean-squared error on urate prediction of 2.147 in the test set. This is also a good predictor of gout, with an AUROC of 0.837 on the validation set (Fig. 15). Allowing pairwise interactions and re-fitting appears to marginally improve urate accuracy, with test error again reducing (from 2.147 to 2.135). As with *LDSmall* and *LDMed*, this more accurate predictor of urate is a worse predictor of gout (AUROC of 0.836).

The training error is significantly smaller than the test or validation error, suggesting there is some overfitting occurring here. The marginal improvement in training, test, and validation loss from interactions does provide some weak evidence of epistasis contributing to serum urate, however.

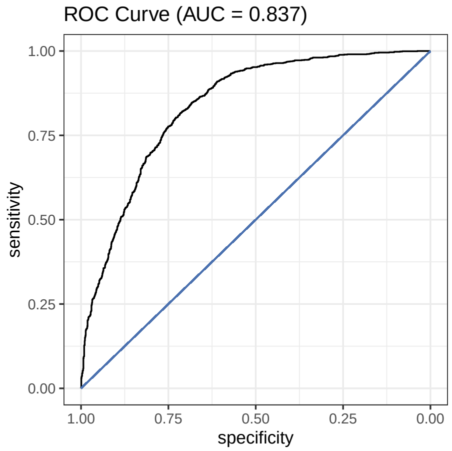


Figure 15: Pint ROC using serum urate predictions in the validation set to classify gout cases. *GWMed* data was used with a limit of $10,000$ non-zero effects and no interaction terms.

**Log Urate** Targeting $log(urate)$ instead of $urate$, we see similar results across all methods (Table 6). The best predictors of $urate$ and $log(urate)$ both achieve AUROC of $\approx 0.837$ when classifying gout, and test loss is typically double the training loss with both $urate$ and $log(urate)$ models. With both $urate$ and $log(urate)$ the best test set urate predictions come from models with interactions fit to *GWMed*, and these models are marginally worse than their interaction-free equivalents at predicting gout.

**Identified Effects** To determine which genes are likely to be responsible for gout, we construct a score for each gene, using only the SNVs with the $1,000$ largest absolute effects $|\beta_i|$ or $|\beta_{i,j}|$. Interaction terms $\beta_{i,j}$ are counted towards both SNVs $i$ and $j$. For each of these SNVs, we check which gene (if any) they are in, then use the sum of these absolute values as the gene's score. We also calculate the mean absolute effect strength assigned to each SNV in the gene. The top effects found in *GWMed* by absolute gene score are shown in Table 7. We see that the overwhelming majority of SNV weight in gene-coding regions is assigned to SLC2A9, a gene known to transport uric acid with variants known to increase risk of gout [3, 35]. ABCG2 is also known to be associate with gout [48], as is SLC16A9 [43]. When fitting *GWMed*, we are restricted to SNVs that are already associated with urate, and finding relevant genes in this case is unsurprising. For comparison we also fit *LDMed* in Table 8. The highest scoring genes in this case have no known association with gout, however. Given that this model is only a marginally worse predictor of gout, it is likely that the SNVs used are correlated with causal variants.

| Gene Name | Sum Strength | Mean Strength |
|---|---|---|
| SLC2A9 | 1.688 | 0.013 |
| SLC2A9/SLC2A9-AS1 | 0.741 | 0.014 |
| ABCG2 | 0.686 | 0.014 |
| RN$F_1$15 | 0.515 | 0.025 |
| SLC16A9 | 0.363 | 0.010 |

Table 7: Top predicted genes in *GWMed* by sum of absolute effect. Note that SNVs in non-coding regions are ignored. Interaction effects were included in this model.

| Gene Name | Sum Strength | Mean Strength |
|---|---|---|
| INS/INS-IGF2 | 0.032 | 0.008 |
| GNPTAB | 0.010 | 0.010 |
| TPH2 | 0.007 | 0.007 |
| PSEN1 | 0.005 | 0.005 |
| CCDC102A | 0.003 | 0.003 |

Table 8: Highest scoring genes in *LDMed*, using sum of absolute SNV effects, including interactions.

If all SNVs were given the same weight, we would score genes that are over-represented in the dataset highly. In the *GWMed* dataset, this makes genes with many SNVs associated with urate particularly likely to score highly, regardless of the predicted SNV effects. To avoid this, we instead take the mean of assigned SNV strengths for each gene. This gives us a different set of top effects, shown in Table 9. Among these, RBM8A, ABCA6, and PKD2 are known to be associated with serum urate or gout [66, 37, 13]. Note that the majority of SNVs were assigned small effects, with a few genes managing significantly higher mean SNV strengths (Fig. 16).

| Gene Name | Sum Strength | Mean Strength |
|---|---|---|
| RBM8A/POLR3GL/LIX1L-AS1 | 0.350 | 0.070 |
| PEX11B/GNRHR2 | 0.041 | 0.041 |
| LINC02806/LINC01731 | 0.039 | 0.039 |
| ABCA6 | 0.079 | 0.026 |
| PKD2 | 0.051 | 0.026 |

Table 9: Highest scoring genes by mean predicted SNV effect in *GWMed*, including interactions.

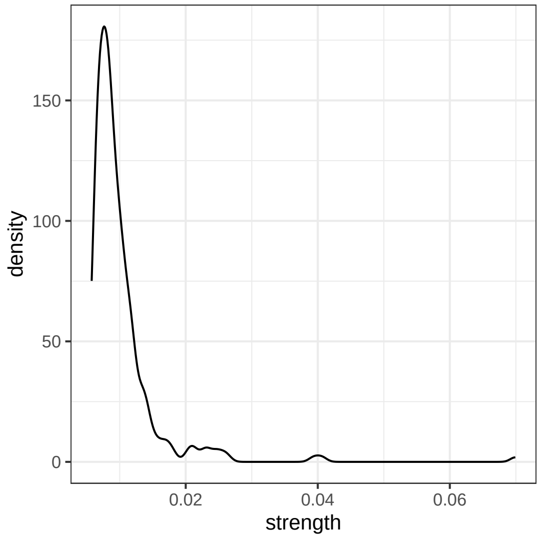


Figure 16: Distribution of mean absolute gene effect strengths, including interactions.

### E Differentiable Logic for UKBB Data

Given that SNVs associated with complex traits are likely to be expression quantitative trait loci (eQTLs) [45], a model that accounts for these regulatory functions may be better equipped to model complex traits. With this in mind, we design a model based on eQTLs, genetic variants that explain some component of the variation in gene expression [44]. We use the same training, testing and verification sets here as for the other methods (Section 5.1). Data is converted into a binary matrix using the method described in Appendix C, with the number of demographic thresholds $k$ left as a tunable hyperparameter.

#### E.1 Expression Quantitative Trait Loci

Typically, hundreds or thousands of loci are associated with the expression of a single gene [57]. Linear models have been able to effectively model many eQTLs [54], and our model therefore assumes that eQTLs have an approximately linear effect on gene expression. We allow for the possibility of a non-linear relationship between gene expression and gout, however.

Our model supposes the expression of each gene $g_i$ is approximately $\beta_0 + \sum_j \beta_j v_j$, where $\beta_j$ is the contribution of the SNV $v_j$. Furthermore, we assume that a complex trait $t$ (gout) occurs when the expression of certain genes are above or below particular thresholds. More formally, $t$ is a proposition that is true when certain gene expression conditions are met. For instance, if gout depends on $g_i$ being greater than $x_i$ and $g_j$ being less than $x_j$, then $t \iff (g_i > x_i) \wedge (g_j < x_j)$, where $x_i$ and $x_j$ are the expression limits of genes $i$ and $j$ respectively. Note that we can approximately model under-expression using negated over-expression, $g_i < x_i \approx \neg(g_i > x_i)$. While this does not account for the case when $g_i = x_i$, in practice we can work around this by choosing a new threshold $x'_i$ such that $g_{obs} < x'_i < x_i$, where $g_{obs}$ is the largest observed value of $g_i$ that is less than $x_i$. We therefore only include lower bounds and negation, but not upper bounds.

Accurately modelling this given only SNVs would require us to identify the contributions of each SNV to both genes $g_i$ and $g_j$, identify the expression limits required for these genes to have an effect, and identify the logical function of these gene effects required for the trait $t$ to be present. While these can be inferred separately, for instance with regression and decision trees, modelling expression separately would require measurements of expression.

We propose instead an end-to-end model that assumes the effects follow this pattern, without specifying the genes involved. Instead we use implied expression values and optimise the entire model directly for complex trait prediction. Our model is based on a linear contribution of SNVs to expression of genes. Whether these expression levels exceed learned thresholds is used as a logical input to a disjunctive normal form expression, which is then used to predict the trait. To optimise this model we need to be able to simultaneously optimise the disjunctive normal form and linear components, as well as represent and choose from a sufficiently wide range of logical formulae.

#### E.2 Disjunctive Normal Form

A logical formula is said to be in disjunctive normal form if it is a disjunction of clauses, each of which is a conjunction of propositions or their negations.

$$(P_1^1 \wedge P_2^1 \wedge \cdots \wedge P_{k_1}^1) \vee (P_1^2 \wedge \cdots \wedge P_{k_2}^2) \vee \ldots (P_1^l \wedge \cdots \wedge P_{k_3}^l)$$

where $\vee$ is the logical 'or' function (disjunction), and $\wedge$ is 'and' (conjunction). Every propositional formula is equivalent to a formula in conjunctive normal form [41]. For simplicity, we assume the logical function for each trait is in conjunctive normal form, e.g.

$$(G_1 \wedge G_2) \vee G_3 \vee (\neg G_4 \wedge G_5)$$

where $G_i$ is true if and only if $g_i > x_i$.

We leave the length of this formula as an adjustable hyperparameter. We further assume that the genes we predict the expression $g_i$ of, are exactly the ones responsible for the trait $t$. Since we make no assumptions about which SNVs contribute to this gene, we can do this without loss of generality.

#### E.3 Model Definition

To simultaneously optimise both the logical and linear components of this model, we define the following differentiable logic, inspired by a many-valued logic defined by Łukasiewicz [34]. In Łukasiewicz logic, we have the following continuous definitions for the 'and', 'or', and 'not' functions:

$$\wedge(a,b) \iff max(0, a+b-1)$$

$$\vee(a,b) \iff min(1, a+b)$$

$$\neg(a) \iff 1-a$$

where $a$, and $b$ are real numbers in the range $[0,1]$, each representing the degree of our belief in some proposition. In our complex-trait model these are the variables $G_i$ indicating whether a gene is over or under expressed. Note that $\wedge$ and $\vee$ are the definitions of strong (rather than weak) disjunction and conjunction. We do not use the implication, equivalence, weak conjunction or weak disjunction operations. Using a real-valued generalisation of classical logic allows us to differentiate the model and learn parameters with gradient-based methods. We need to be able to not only differentiate the correct logical function, but to choose what that function should be. For the latter, we introduce a *weighted* version of Łukasiewicz logic.

$$\begin{aligned} \wedge(a,b,w_a,w_b) &\iff max(0, w_a \cdot a + w_b \cdot b - 1) \\ \vee(a,b,w_a,w_b) &\iff min(1, w_a \cdot a + w_b \cdot b) \\ \neg(a) &\iff 1-a \end{aligned} \tag{9}$$

The weights $w_a$ and $w_b$ change the extent to which $a$ and $b$ are included in the 'and' or 'or' functions. Using these weights, we can extend these functions to many-valued functions by replacing $a$ and $b$ with an arbitrarily long vector $a$:

$$\begin{aligned} \wedge(a, w) &\iff max(0, 2 \cdot a \cdot softmax(w) - 1) \\ \vee(a, w) &\iff min(1, 2 \cdot a \cdot softmax(w)) \end{aligned} \tag{10}$$

where $softmax(x)$ is the function $softmax(x_i) = \frac{e^{x_i}}{\sum_j e^{x_j}}$. This allows us to use weights as a gradually adjustable form of variable selection. Finally, to ensure the function is differentiable outside the range $[0, 1]$, we replace the $min$, and $max$ hard thresholds with the function $r$ (Eq. (11) and Fig. 17).

$$r(x) = \begin{cases} e^{x+log(0.2)-0.2} & \text{if } x < 0.2 \\ 1.0 - e^{1.0-x+log(0.2)-0.2} & \text{if } x > 0.8 \\ x & \text{otherwise} \end{cases} \tag{11}$$

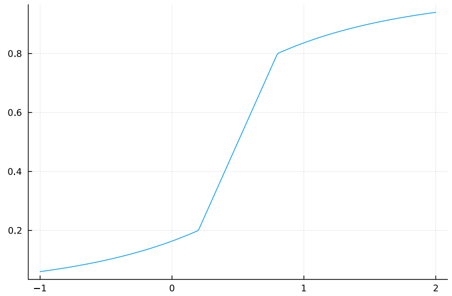


Figure 17: $y = r(x)$ as defined in Eq. (11).

The final 'and' and 'or' functions are differentiable at all points, and outputs remain within the range $(0, 1)$:

$$\begin{aligned} \wedge(a, w) &\iff r(2 \cdot a \cdot \mathrm{softmax}(w) - 1) \\ \vee(a, w) &\iff r(2 \cdot a \cdot \mathrm{softmax}(w)) \end{aligned} \tag{12}$$

Using these operations, we define the model as follows. Expression values $e$ are a linear combination of SNV values $X_{snv}$, using weights $W_l \in \mathbb{R}^{n \times l_{out}}, W_{l,0}$, where $n$ is the number of SNVs, and $l_{out}$ is a hyperparameter deciding how many genes we are estimating the expression of.

$$e = W_l X_{snv} + W_{l,0}$$

For each demographic value $d_i$ (age and BMI) we replace the variable $d_i \in \mathbb{R}$ with $k$ binary values and combine this with the SNV matrix using the method described in Appendix C. This gives us a binary demographic vector $D$. We concatenate the expression and demographic vectors into a combined binary vector $X_c = [D, e]$. Next, we get the matrix $X'$ with each value and its negation $X' = [X_c, \neg X_c]$. We then get the values of the $c$ conjunctions $a_1, \ldots, a_c$ using variable selection weights $w_1^{and}, \ldots w_c^{and}$:

$$a_i = \wedge(X', w_i^{and})$$

The final output is that of the disjunction, with variable selection weights $w^{or}$:

$$\vee(a, w^{or})$$

The complete model has learned parameters $w_1^{and}, \ldots w_c^{and}$ for each of the $c$ conjunctions, $w^{or}$ for the disjunction, and $W_l$ for the linear function of SNVs. We implement our model using Lux [47] and MLJ [6], using the AdamW algorithm to optimise parameters [33].

**Binarised Model** When two weights $w_i$ and $w_j$ in $w$ are much larger than all others, $2 \cdot softmax(w)$ gives near zero weight to other inputs, while $i$ and $j$ will have weight near one if $w_i \approx w_j$. With extreme weights, behaviour therefore approaches that of a binary model. This binary model has the classical logic 'and' and 'or' functions, and is easier to interpret than the weighted model. To convert a trained weighted model to a strictly binary one, we convert the weights from $w \in \mathbb{R}^n$ to $w' \in \{0, 1\}^n$ as follows.

$$w'_i \leftarrow (w_i > 0)$$

If $w'_i = 0$ for all $i$, then we assign 1 to the index with the largest value: $\max_i w_i \leftarrow 1$. These are the weights that have increased, rather than decreased, which we interpret as meaning they should be included in the expression. Using these new weights $w'$, we revert to the hard $[0, 1]$ limit functions in Eq. (9) and use the new weights directly without the softmax operation (Eq. (13)).

$$\begin{aligned} \wedge(a, w') &\iff min(1, max(0, a \cdot w' - (\|a\|_1 - 1))) \\ \vee(a, w') &\iff min(1, max(0, a \cdot w')) \end{aligned} \tag{13}$$

This allows us to learn parameters in the continuous model using gradient descent, then convert the learned continuous model to a classic binary model if the weights converge to extreme values.

#### E.4 Hyperparameter Optimisation

As with LightGBM, we run a hyperparameter search using Latin Hypercube sampling [2] with two generations and a population size of 120, implemented in MLJ [5]. The following four hyperparameters are optimised: the number of SNVs in each linear eQTL function, for which we test values between 1 and 32; the number of eQTL gene regulation functions, using the same range; the learning rate, varying from $10^{-7}$ to $10^{-1}$ on a log scale; and the number of binary variables used to represent each demographic value, varying between 1 and 20. We encode demographic information using the same binary matrix as in Appendix C. The model is run for 20 epochs, using the training data described in Section 5.1 to predict gout. The results of this hyperparameter search, including test set cross-entropy, are summarised in Fig. 18. Hyperparameter trials were performed on an and AMD EPYC 7713 64-Core Processor, on a system with 2TB of memory.

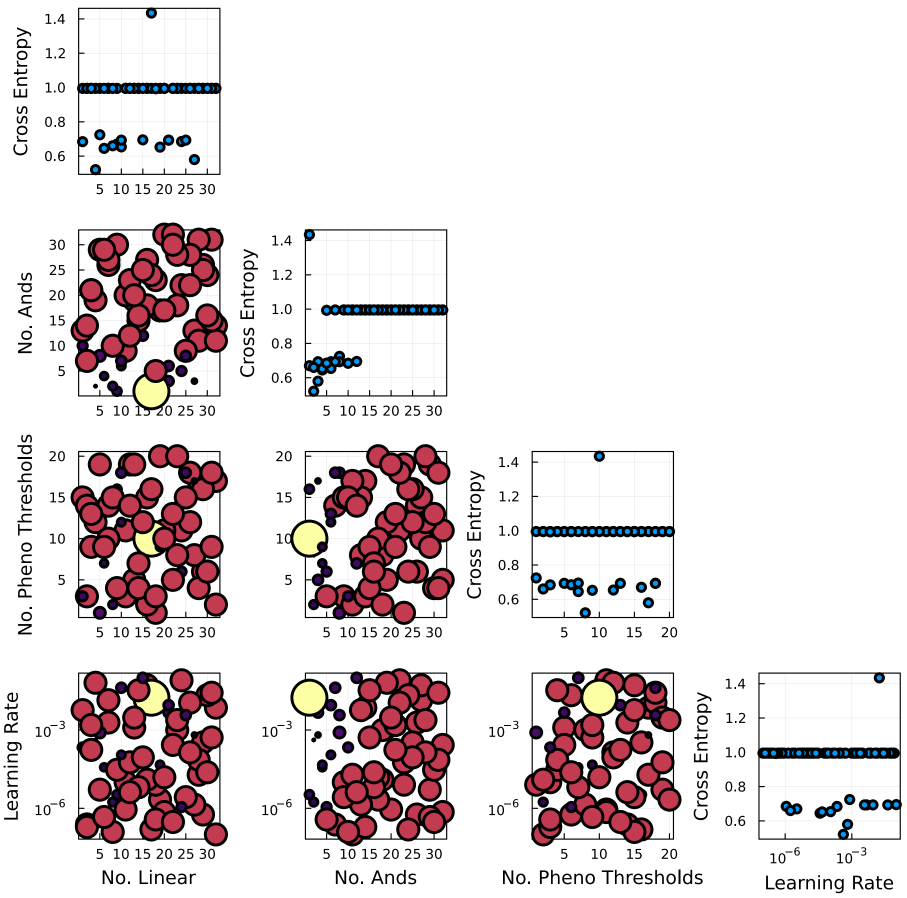


Figure 18: Logic model hyperparameter search. Test-set cross entropy with respect to each parameter is shown on the diagonal, while relative hyperparameter distributions are shown below. *No. Linear* is the number of linear gene expression functions modelled. *No. Ands* is the number of conjunction clauses in the disjunctive normal form formula. *No. Pheno* Thresholds is the number of binary variables used to represent each demographic value. Point size and color in parameter vs. parameter plots indicate test-set loss of the trial, with smaller darker points being more successful (lower loss).

The one parameter with a clear impact is the number of 'and' terms included in the final disjunction, with models that had few 'and' terms performing significantly better in terms of test set cross-entropy (Fig. 18). This parameter determines the number of conjunctive clauses in the disjunctive normal form. Using a small number of conjunctive clauses limits the complexity of the logical function, which in the extreme case (one linear function and one conjunctive clause) reduces to linear regression. The best performing model had 4 linear functions, 2 conjunctive clauses, 8 demographic thresholds, and a learning rate of $4.160 \times 10^{-4}$. This model has an AUROC of 0.790 in the validation set (Fig. 19), behind both logistic (Appendix C) and linear (Appendix D) regression. We construct binary versions of the best model for each dataset using the method in Appendix E.3, however these are no better than random (AUROC of 0.5) at classifying gout.

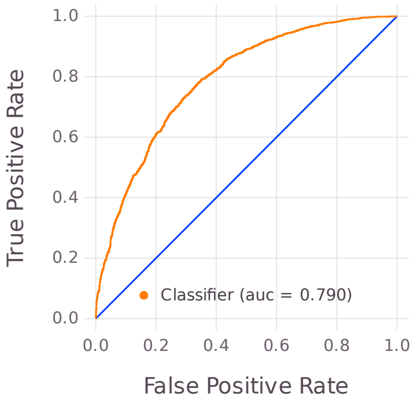


Figure 19: Best performing logic model ROC curve in the validation set.

### F Transformer Variations

Deep neural networks, including transformer-based networks initially designed for natural language processing (NLP) tasks [70], have recently been successfully employed to analyse molecular sequence data, the most successful and widely-known example being AlphaFold-2 [25].

Presently, however, such architectures are capable only of handling sequences of modest length, such as those encoding specific proteins. Recently, and in parallel, transformer-based architectures for NLP have been developed that aim to accommodate extended bodies of text [72, 7]. Here, we combine these two directions of research and present a transformer-based deep neural architecture for GWAS data, including a purpose-designed SNV encoder, that is capable of modelling gene-gene interactions and multidimensional phenotypes, and which scales to a significant fraction the whole-genome sequencing data standard for modern GWAS. Given their ability to encode complex patterns in text sequences, transformer networks have the potential to capture complex relationships between SNVs. By doing so, they may be able to outperform classic approaches in complex trait prediction.

We begin by using a simple encoder architecture to classify gout cases, demonstrating the potential of transformer networks. We do not perform any hyperparameter optimisation with this encoder, and test it using 25% of the gout cases and a matching number of non-gout cases. Expanding on this, we test a number of variations on the input encoding, network architecture, and hyperparameters. We describe each of these variations, comparing the cross-entropy loss in the test set predictions with each option. For each choice of hyperparameters, we train using 70% of the gout cases and a matching number of non-gout cases. The test and validation sets are balanced sets containing 15% of the gout cases each. We use the same test set to evaluate each choice of hyperparameters, reserving the validation set for the final analysis. We refer the training and testing of a set of hyperparameters as a trial.

#### F.1 SNV Encoding

Each SNV can be defined by two nucleotide sequences, a reference value and a possible alternative, and their position in the genome. For example, an SNV might be at position 10, the 10th nucleotide of the genome, where the reference genome might contain an 'A', but in some cases an insertion changes this to 'AG'. We call the two candidates at a given position alleles. It is important to note that every chromosome in the human genome is present twice, and there may be differences between these two copies. For a given SNV, individuals then have either two copies of the first allele, two of the second, or one of each. Note that we do not consider the case where more than two possibilities exist at the same position.

To use this information in a neural network it must be encoded into a vector. We do this by first converting each case of each SNV into a token, then using learned fixed-length vectors ('embeddings') to represent these tokens (see Appendix F.3 for details). Note that self-attention is based on the dot-product similarity of encoded vectors, meaning the similarity of learned vectors affects the self-attention scores.

We test several variations of the encoding. These encodings represent SNVs as integer tokens while preserving different information about the differences between the two alleles. Except for version 1 (below), these all remove some information to avoid long insertions or deletions being encoded simply as unique changes. While encoding every unique variation as a unique token allows us to distinguish every change present in the data, it also ignores any similarity that might be present. By choosing exactly which information is included in the encoding we decide which differences should be ignored, and make associations between otherwise similar variants easier to detect. This is done by reducing long insertions or deletions to shorter sequences in a variety of ways, resulting in identical shorter sequences in similar cases (e.g. all long insertions after a G will be treated as 'insertion after G' in encoding V2).

Regardless of the choice of encoding, we encode the following additional special tokens. When it is unknown which alleles of an SNV are present, we encode this as 'nan'. We reserve 'cls' for the classification token, and 'mask' for masking input sequences. Each token in the dataset is represented with a unique integer. With tokens represented by integers $i \in \{1, \ldots, k\}$, we then use these integers as indices to determine which learned embedding vector $e_i \in \mathbb{R}^d$

is passed to the neural network. The encoding process is shown in Fig. 20.

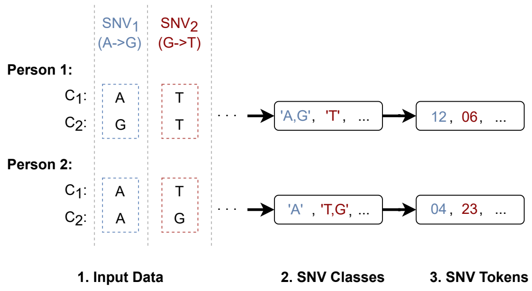


Figure 20: Pre-processing of SNV sequences. Given an input set of SNVs for each person, the combination of SNVs and zygosity is mapped to one of several cases. Each case is encoded as an integer, and each person's sequence is encoded as a sequence of these integers. This example uses encoding V1.

**V1** Encoding V1 creates a string of characters out of the nucleotide sequence present at the position of an SNV, combining both sequences when two different alleles are present. Each unique string encountered in the dataset is then represented with a unique integer. Let the SNV be defined by two nucleotide sequences $a$ and $b$ at some position. If in both alleles we have the sequence $a$, then our string is simply '$a$'. Similarly, if both alleles contain $b$ then we represent simply '$b$'. When one allele contains $a$ and the other $b$, we use the string '$a, b$'. For example, if the SNV described an insertion of 'G' after the nucleotide 'A' at some position, and one of each allele was present, we would represent this as the string 'A,AG'. If this were the first SNV encountered in the data, it, along with all other instances of 'A,AG', would be encoded as 1. Under this encoding, long unique sequence variations, of which there are many, will be represented as unique tokens.

**V2** Encoding V2 resembles encoding V1, but attempts to limit the number of unique tokens. This is inspired by the intuition that, in many cases, it is not the exact sequence of a long insertion that is important, but rather the fact that an insertion was present. We therefore represent all insertions using a common character 'I', rather than their (likely unique) sequences. Specifically, we cut of all strings after the first character, then add an 'I' if other characters were present. For example, 'GTTG' becomes 'GI'. This reduces long sequences down to only a few possible combinations.

When both alleles are then represented by the same string $a$, we encode this as '$a$' as before. When one of each of the variations $a$ and $b$ is present, but both are the same length, we again encode this as '$a, b$'. To explicitly distinguish insertions and deletions, when $b$ is longer than $a$ we encode this as '$a, ins$'. Similarly, when $b$ is shorter than $a$ we encode this as '$a, del$'. Encoding *GWLarge* in this way gives us 33 unique tokens, shown in Table 10.

| | | | | |
|---|---|---|---|---|
| ‘nan’ : 00 | ‘ins’ : 01 | ‘del’ : 02 | ‘G’ : 03 | ‘A’ : 04 |
| ‘C’ : 05 | ‘T’ : 06 | ‘CI’ : 07 | ‘GI’ : 08 | ‘TI’ : 09 |
| ‘AI’ : 10 | ‘G,A’ : 11 | ‘A,G’ : 12 | ‘G,C’ : 13 | ‘C,T’ : 14 |
| ‘G,T’ : 15 | ‘C,G’ : 16 | ‘T,C’ : 17 | ‘A,ins’ : 18 | ‘A,C’ : 19 |
| ‘CI,del’ : 20 | ‘G,ins’ : 21 | ‘GI,del’ : 22 | ‘T,G’ : 23 | ‘C,A’ : 24 |
| ‘TI,del’ : 25 | ‘A,T’ : 26 | ‘C,ins’ : 27 | ‘T,ins’ : 28 | ‘AI,del’ : 29 |
| ‘T,A’ : 30 | ‘AI,ins’ : 31 | ‘cls’ : 32 | ‘mask’ : 33 | |

Table 10: SNV encoding V2. Homozygous alleles are encoded as ‘X’, heterozygous as ‘X,Y’. ‘I’ encodes all nucleotides after the first.

**V3** Encoding V3 is identical to encoding V2, with one exception. When $b$ is longer than $a$, instead of encoding this as ‘$a, ins$’, we encode this as ‘$a, b$’. This distinguishes cases where there is strictly an insertion, e.g. ‘G, GI’, from cases where there is both a substitution and an insertion, e.g. ‘G,TI’.

**V4** This encoding is designed to establish whether the sequence length has changed, and whether it has become longer or shorter. We distinguish the following cases:

- A single nucleotide substitution, A for B, is encoded as ‘A’, ‘B’, or ‘A,B’, as in encoding V1.
- If $a$ is longer than $b$, this is encoded as ‘longer’ if two copies of $a$ are present, ‘shorter’ of two copies of $b$ are present, or ‘mixed_indel’ if one of each is present.
- If $b$ is longer than $a$, this is encoded as ‘longer’ if two copies of $b$ are present, ‘shorter’ for two copies of $a$, and ‘mixed_indel’ for one of each.
- When $a$ and $b$ are both longer than one nucleotide, but the same length, this is encoded as ‘long_sub’ in case two copies of either $a$ or $b$ are present, and ‘mixed_long_sub’ of one of each is present.

**V5** In encoding V5 we ignore the sequences $a$ and $b$ entirely, and simply encode whether two copies of $a$ were present, one of each, or two copies of $b$. These are encoded as ‘0’, ‘1’, and ‘2’, respectively. Note that these are the encoded strings, not the integer values. Since special tokens (‘nan’, ‘cls’, and ‘mask’) are included first, the actual integer values will be larger.

##### F.1.1 Chromosomes and Positions

For our network to correctly identify interactions, the encoding of positions of SNVs in the genome is essential. Broadly speaking, there are two approaches to positional encoding in transformer networks, direct representation and learned, where the direct representation can be of the position in the sequence, or the position relative to other SNVs [15]. We implement and test both sine/cosine direct encodings used in [71], and learned position embeddings, where a separate learned embedding is used for each possible position. Chromosome and SNV positions are encoded separately using the same type of positional encoding.

Effectively, the sine/cosine position encodings give attention to nearby positions in the sequence. While later layers in deep networks can use this information in different ways, initially giving higher attention to nearby SNVs would make sense if nearby SNVs are likely to be related.

To determine whether this is the case, we calculate the pairwise correlation $c_{i,j}$ between every pair of SNVs $i$ and $j$ in the unimputed dataset[1]. We then calculate the mean correlation between SNVs at a given offset by averaging the absolute correlation between all SNVs a fixed distance apart $m_x = \sum_{i=1}^{p} |c_{i,i+x}|$. The mean correlation between every SNV and its immediate predecessor for example is $m_{-1}$. We group these offsets into batches of 50, and plot the distribution of correlation scores by relative position (Fig. 21), excluding $m_0$ from the plot. We see that the position is an excellent indication of the correlation between two SNVs.

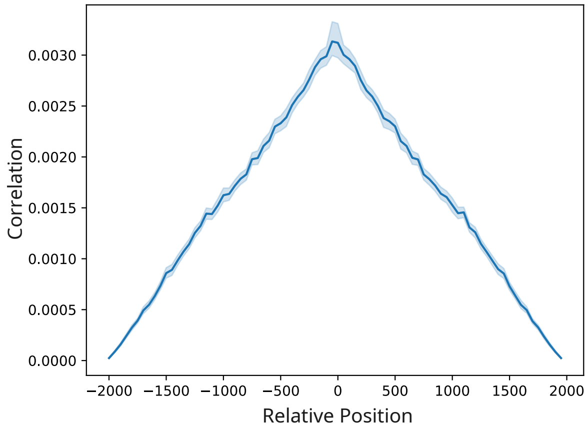


Figure 21: Correlation between SNVs by relative position in sequence. Position is the relative position of the batch of 50 indices.

#### F.2 More General Encoder-Only Architecture

Our transformers, like the one in [18] are based on an encoder-only transformer architecture (as opposed to an encoder-decoder architecture) [71], using Linformer self-attention [72]. The encoder inputs and outputs matrices $X \in R^{n \times d_{embed}}$, where $n$ is the number of SNVs in the dataset, and $d_{embed}$ is a tunable embedding size.

[1] We restrict our analysis to the unimputed dataset because using the full dataset would be computationally challenging.

For classification we place a unique classification token at the beginning of the SNV sequence, then use a single fully connected layer from the final embedding of the classification token to each class. Finally, the softmax function ensures the outputs sum to 1 to form a probability distribution.

**Multi-head Attention** Every attention function consists of several layers. First, since we use Linformer, we project the keys and queries into a smaller dimensional space with $E \in \mathbb{R}^{n \times k_l}$ and $F \in \mathbb{R}^{n \times k_l}$. We then use the scaled-dot-product multi-head self-attention function to produce weighted outputs $O_1 \in \mathbb{R}^{n \times d}$. As we will discuss in Appendix F.6, many of these parameters are tunable.

#### F.3 Vectoriser

To allow testing of various input encodings and architectural changes, we place an extra layer before the encoder that we call the vectoriser. The vectoriser is responsible for all aspects of the input encoding, and can be configured in a number of ways. The overall network then has two components, the vectoriser (Fig. 22) and the encoder. The vectoriser encodes the relevant information for each SNV according to its parameters, using one of the encodings in Appendix F.1, as well as optionally including the SNV's chromosome, position within the chromosome, and gene (if applicable). This encoded matrix is then used as the input to the transformer encoder. The vectoriser takes the input SNV sequence and converts it into an $\mathbb{R}^{n \times d_{embed}}$ matrix, where $n$ is the length of the sequence and $d_{embed}$ is the total embedding dimension.

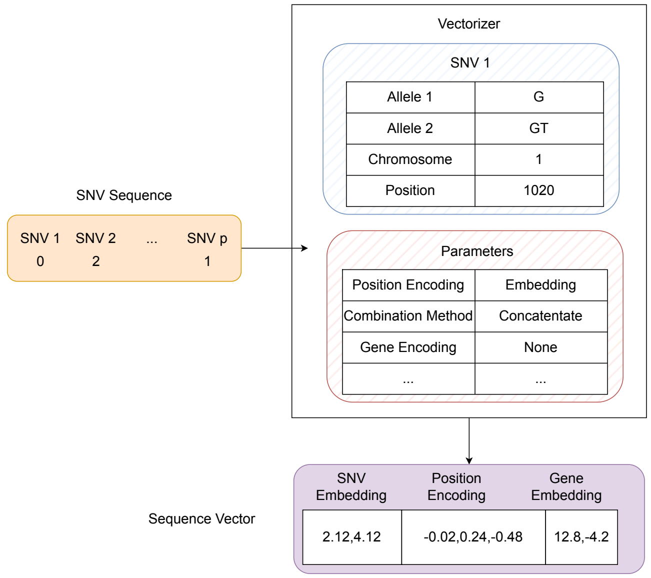


Figure 22: SNVs flow through the vectoriser to be encoded as vectors.

#### F.4 Linformer

While we could use only small subsets of SNVs, the *GWSmall* or *LDSmall* datasets for example, our goal is to include every SNV that could be involved in gout. To that end, we also consider larger datasets, with $\approx 13,000$ SNVS (*GWMed*, *LDMed*), and $\approx 65,000$ SNVs (*GWLarge*). The original transformer self-attention operation $softmax(\frac{QK^{\intercal}}{\sqrt{d}})V$ requires calculating the $n \times n$ matrix $QK^T$, and has time complexity of $O(n^2d)$. As a result it does not scale to tens of thousands of tokens. To overcome this, we use the Linformer self-attention mechanism from Wang et al. [72], internally projecting the $\mathbb{R}^{n\times d}$ sequence representation into a smaller $\mathbb{R}^{n\times k_l}$ matrix for some operations. Note that $k_l$ is a tunable parameter that can be used to balance the representational power of the network with its computational cost.

#### F.5 Hyena

Instead of Linformer, we could use Hyena attention [51]. Hyena is a recent subquadratic attention mechanism that has been used in transformers for genetics [42]. To determine whether we should focus on Hyena or Linformer networks, we train two otherwise equivalent networks on the *LDMed* reduced dataset for 50 epochs. Both networks have 4 layers, an embedding dimension of 64, and 4 attention heads. We use a Linformer $k$ value of 64 for testing. The ROC curve in the validation set for each method is shown in Fig. 23.

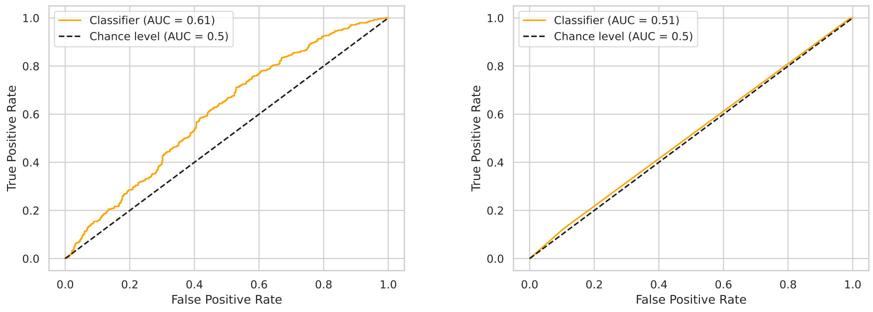


(a) (b)

Figure 23: Linformer (a) vs. Hyena (b) encoder with equivalent parameters. Test is performed on the test set after 50 training epochs on *LDMed* dataset.

For our purposes, it appears Linformer significantly outperforms Hyena attention (Fig. 23), with AUROC of 0.61 vs. 0.51. While it may be possible to develop a Hyena-based network with comparable performance, we focus our attention on the faster Linformer approach.

#### F.6 Hyperparameter Search

Beginning with the architecture described in [18], we developed a number of optional variations. To choose an effective combination of these options, as well as appropriate hyperparameters, we use the AHSA scheduler [31] implemented in the Ray Tune hyperparameter tuning library [55].

As well as the basic tunable parameters, embedding dimensions, number of heads, network depth, learning rate, batch size, and dropout, we test several more substantial architectural changes in our hyperparameter search:

- Including gene and chromosome embeddings.
- Embedding a graph of known gene interactions.
- Alternative SNV encodings.
- Alternative weight initialisation methods.
- Alternative position encodings methods.
- An alternative output classification block.

**Gene and Chromosome Embedding** For every SNV, we optionally include a learned embedding of its gene and chromosome, if applicable. This allows us to identify relationships between genes, and associate SNVs in the same gene or chromosome. Particularly in pre-training, this should lead to higher attention between SNVs in correlated genes. We do not need to learn these relationships from scratch, however.

We either use a simple learned embedding, or optionally encode genes using a Graph Convolutional Network [29, 20], with a node for each gene present in the dataset. Edges are placed between nodes when an interaction of any kind between the corresponding genes is present in the BioGRID interaction database [46]. Given the graph adjacency matrix $A$, input gene embedding $X$, and learnable weight parameters $W_0$ and $W_1$, the graph network outputs a vector $Z \in \mathbb{R}^{n \times d}$ where the embedding of every gene is influenced by all genes connected in the graph.

$$\begin{aligned} \tilde{A} &= A + I_N \\ \tilde{D}_{ii} &= \sum j \tilde{A}_{ij} \\ \hat{A} &= \tilde{D}^{-\frac{1}{2}} \tilde{A} \tilde{D}^{-\frac{1}{2}} \\ Z &= d(\hat{A}\ relu(\hat{A} X W_0) W_1) \end{aligned} \tag{14}$$

We use the same embedding dimension $d_{embed\_genes}$ for nodes in the graph network and hidden layers. These output embeddings retain the input dimensions $n_{genes} \times d_{embed\_genes}$, and are used instead of the randomly-initialised learned embeddings of the simple gene encoding above. The complete SNV encoding with gene-graph encoding is shown in Fig. 24.

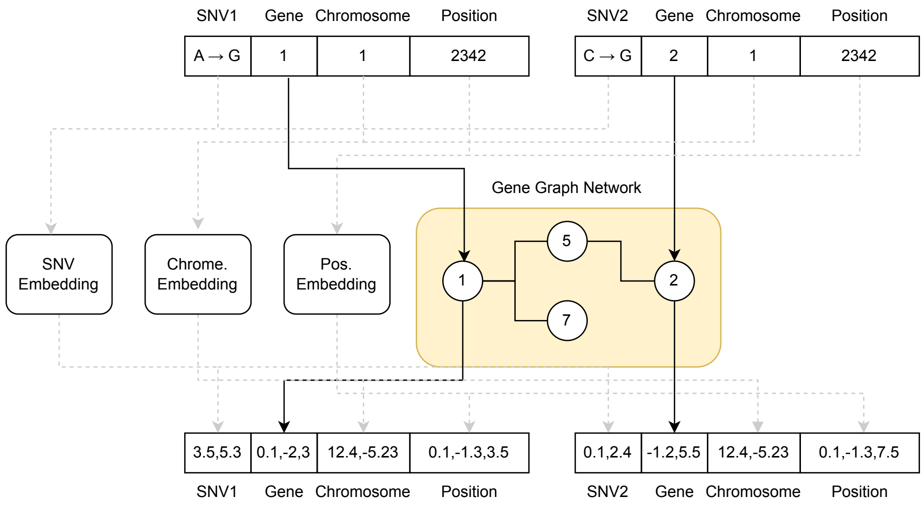


Figure 24: SNV embedding with graph neural network gene encoding.

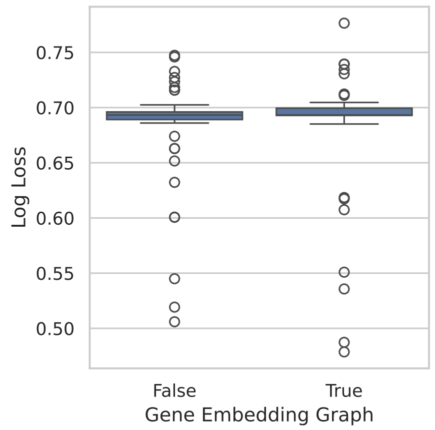


Figure 25: Gene graph used vs. loss. False implies the gene graph was not used, which includes cases where no gene embedding was included.

While there is no significant advantage on average, the two best-performing networks did use the gene graph (Fig. 25), suggesting that with the right parameters this model can improve performance.

**Encoding Version** All of our encoding variations distinguish SNVs present in one allele, two alleles, or not at all. The only difference is whether or how they distinguish different types of variation, i.e. insertions, deletions, or specific substitutions. To determine whether there is any significant difference between them, we fit the best-performing Linformer network from our automatic hyperparameter search to the *LDMed* dataset using each encoding method. Note that this network was initially chosen using a hyperparameter search on data with encoding version 5 (see Appendix F.7).

As we see in Table 11, there is almost no difference in performance when using different encoding versions. While $V1$ does marginally outperform $V5$ here, the difference was not reproducible. Since the difference between $V1$ and $V5$ is therefore less than the variance between repeated runs with $V5$, and we cannot conclude that either is generally preferable. We therefore conclude that distinguishing specific substitutions, insertions, or deletions, is not necessary, and use encoding V5 in the remainder of this chapter. This has the advantage that, since there are relatively few unique tokens, encoding these tokens may be more efficient.

| Encoding Version | AUROC |
|---|---|
| V1 | 0.83 |
| V2 | 0.81 |
| V3 | 0.81 |
| V4 | 0.82 |
| V5 | 0.82 |

Table 11: Comparison of AUROC given different input encodings.

**Dropout** Most layers of the transformer network have a dropout parameter, which randomly zeroes some elements of the input with probability $p$ using samples from a Bernoulli distribution. The dropout parameter and corresponding cross-entropy loss for each trial in the hyperparameter search are shown in Fig. 26. All trials achieving test set loss below 0.6 had dropout near 0.1, although this is also where we see the majority of samples (Fig. 26).

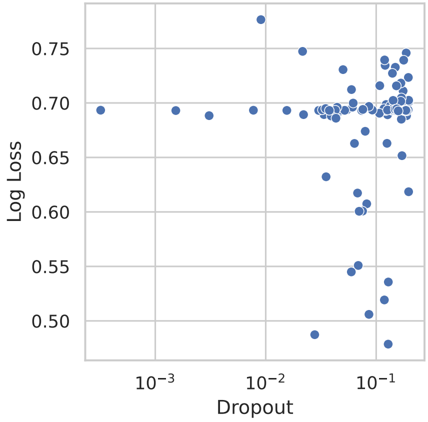


Figure 26: Dropout vs loss. Each point represents a separate hyperparameter trial. Note that other hyperparameters also vary between trials.

**Number of Attention Heads** As described in Appendix F.2, we use multiple attention heads, following Vaswani et al. [71]. This divides the embedding dimension $d_{embed}$ among $n_h$ heads, so that each provides separate scores based on an embedding of size $\frac{d_{embed}}{n_h}$. In our hyperparameter optimisation we sample $n_h$ uniformly at random from $\{1, 2, 4, 8\}$. If the chosen embedding dimension $d_{embed}$ cannot be divided by $n_h$, we increase the size of the SNV embedding $d_{embed-snv}$, and therefore the total embedding size, by $n_h - (d_{embed} \mod n_h)$.

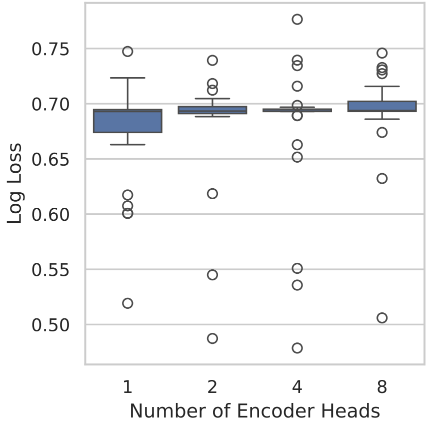


Figure 27: Number of attention heads vs. test set gout prediction cross-entropy loss.

A smaller number of heads appears advantageous on average, with two or four heads producing the best performing networks. There is, however, no clear significant difference between one, two, four, or eight heads (Fig. 27).

**Feed-Forward Network Scale** The feed-forward network dimension $d_f$ can be any natural number. In practice it is often set to a multiple of the input vector length $n$. We set $d_f$ to be a multiple $k_f$ of $n$, and sample $k_f$ uniformly at random from $\{1, 2, 4, 8\}$ in our hyperparameter search.

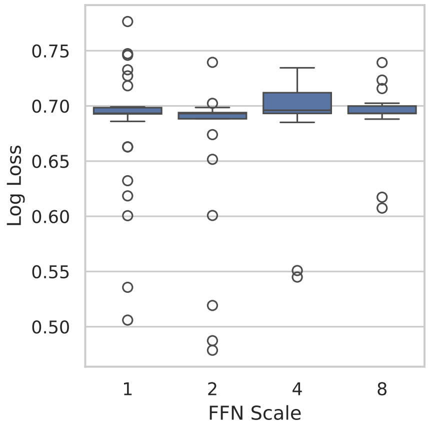


Figure 28: Feed-forward network hidden layer scale vs. test set loss.

Transformer-block feed-forward networks two times larger than the input size produced the two best-performing trials, although the median performance was similar across all scales (Fig. 28).

**Embedding Dimension** The overall embedding dimension is determined by the SNV and gene embedding dimensions $d_{snv}$ and $d_{gene}$, as well as the chromosome and position embedding dimensions $d_{chom}$ and $d_{pos}$, and, depending on the choice of hyperparameters, the number of attention heads.

$$d_{embed} = d_{snv} + d_{gene} + d_{chrom} + d_{pos} + d_{offset}$$

We exclude $d_{chrom}$ if the chromosome is ignored, and $d_{pos}$ if the position is being added rather than concatenated. To ensure the number of heads is a factor of the total embedding dimension we add $d_{offset} = n_{heads} - (d'_{embed} \mod n_{heads})$, where $d'_{embed}$ is the value of $d_{embed}$ if the offset were zero. Instead of optimising the overall embedding size, we optimise each of these smaller embeddings on their own.

**SNV Embedding Size** SNV embeddings are learned embeddings, one for each possible SNV token. For the hyperparameter search we sample the size of these embeddings uniformly from $\{2, 4, 8, 16\}$.

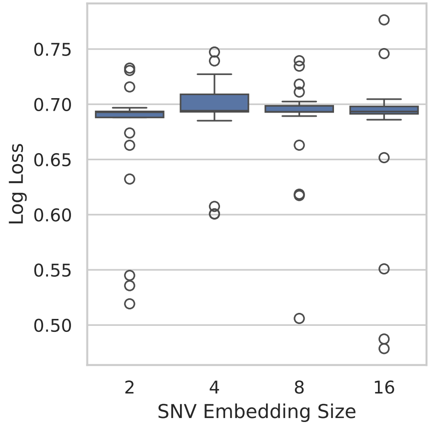


Figure 29: SNV embedding size vs. test set loss.

There is no clear advantage to any particular SNV embedding size, although embeddings of size 16 were used in the two top performing trials (Fig. 29).

**Gene Embedding Size** For each SNV, we look up its position in the human genome assembly GRCh37 [9] and determine whether it is positioned inside a gene. We maintain a fixed-size learned embedding for each gene that has at least one SNV in our sequence, and include this in our SNV embedding. SNVs outside a gene all get the same (learnable) embedding. We sample the gene embedding size uniformly from $\{0, 4, 8, 16, 32\}$. For size 0, the gene embedding is not used.

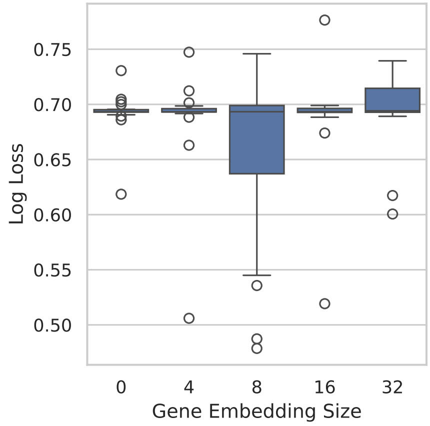


Figure 30: Gene embedding size vs. test set loss.

A gene embedding of size 8 clearly outperforms both smaller and larger choices (Fig. 30).

**Chromosome Embedding** For each SNV, we either include the chromosome as a fixed-size learned embedding, or ignore it entirely. When the chromosome embedding is included, we sample its size uniformly from $\{8, 16, 32\}$.

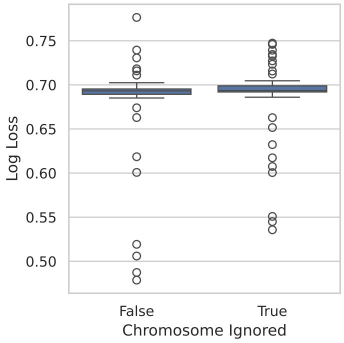


Figure 31: SNV Chromosome ignored vs. test set loss.

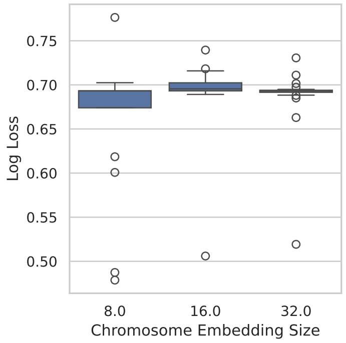


Figure 32: Chromosome embedding size vs test set loss.

The four best performing networks all included the chromosome, although there is no clear advantage on average (Fig. 31). Among trials using the embedding, we see a noticeable improvement in performance using a smaller chromosome embedding size (Fig. 32).

**Position Encoding** The vectoriser (Appendix F.3) can encode the position in one of two ways. With the sine/cosine encoding, position embeddings are produced with alternating sine/cosine functions of the position at increasing frequencies. Note that the majority of positions in the genome do not have a corresponding SNV in our data, and as a result genome positions are not the same as indices in the sequence.

Alternatively, we use a learned embedding, in which position embeddings are a learned vector, one per position in the sequence. These are initialised to the sine/cosine encoding of the sequence $1, \ldots, n$ for a sequence of length n, but are then learned as a standard parameter.

In either case, we sample the embedding vector size uniformly from $\{8, 16, 32\}$.

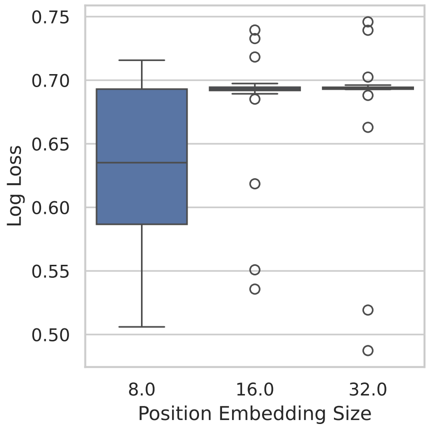


Figure 33: Position embeding size vs. test set loss.

Smaller position embeddings had a significant advantage over larger ones on average, although outliers performed comparably well for all sizes (Fig. 33).

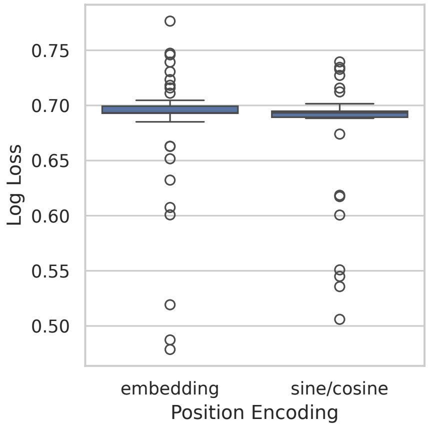


Figure 34: Position encoding method vs. test set loss.

There is no consistent advantage between embedding and sine/cosine position embeddings, although the two best performing networks used learned embeddings (Fig. 34).

**Position Embedding Combination Method** The encoded position can be included in the complete SNV embedding either by concatenating with the SNV (and optionally other) encoding(s), $e = [e_{SNV}, e_{pos}]$, or by adding it, $e = e_{SNV} + e_{pos}$.

Concatenating increases the embedding dimension, and therefore the number of parameters required for the encoder, significantly. It has the advantage, however, of allowing the position encoding to be used without any risk of conflating the SNV and position encodings. We use either method with a 50% chance in our hyperparameter search.

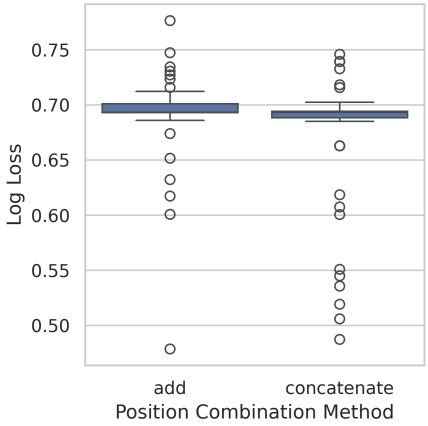


Figure 35: config pos combine vs loss

Both high and low performing tails showed a general advantage to concatenating, rather than adding, the positional encoding to the rest of the SNV embedding (Fig. 35). The single best performing network is using addition however, suggesting there is no consistent advantage.

**Number of encoder Layers** Theoretically we can have any number of encoder layers $n_l$. In practice, however, we quickly run out of GPU memory when $n_l$ is large. In our hyperparameter search we sample $n_l$ uniformly from $\{1, 2, 3, 4, 5, 6, 7\}$.

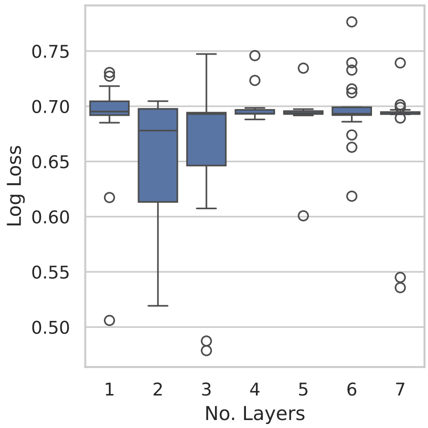


Figure 36: Number of encoder layers vs. test set loss.

We see a clear advantage to networks with two or three layers, with significantly better average performance than other options. There were also outliers with good performance using one or seven layers (Fig. 36).

**Batch Size** We use batches largely to improve parallelism. Averaging the gradients of a batch of inputs does have an effect on the final parameters however, since we use the same initial parameters for each item in the batch rather than adjusting them and re-calculating the gradient each time. We sample batch sizes uniformly from $\{1, 2, 4\}$.

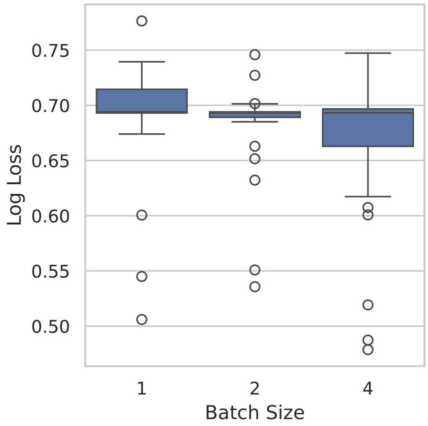


Figure 37: Batch size vs. test set loss.

We generally see larger batch sizes performing better (Fig. 37), and batches larger than four samples might improve performance further. Larger batches require increasing amounts of memory, however, and we cannot increase the batch size significantly beyond four without exhausting GPU memory.

**Linformer K** The Linformer projection significantly reduces the size of our query and key vectors for the sake of performance. Lower values of $k_l$ reduce the memory requirement and running time, however they risk discarding too much information. Larger values of $k_l$ allow the model to encode more information about the sequence at the cost of time and memory. We sample $k_l$ uniformly at random from $\{16, 32, 64, 128, 256\}$.

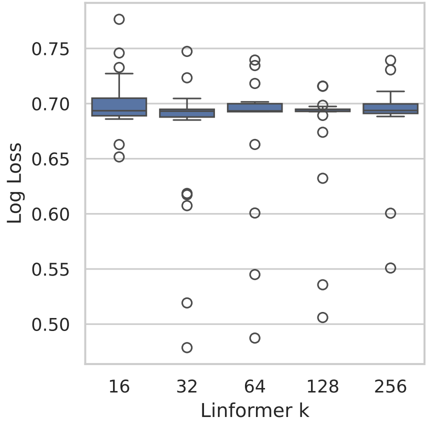


Figure 38: Linformer k vs. test set loss.

There is a clear disadvantage to a Linformer k value of only 16. There is no consistent winner among the larger values, however, although the two best performing trials had values of 32 and 64 (Fig. 38).

**Ignore Class** We optionally include a one-hot encoding of the type of token we include at each position, which is either a class token 'cls', a mask token 'mask', an SNV, or a demographic value. We have a 50% chance of instead ignoring the class in our hyperparameter search.

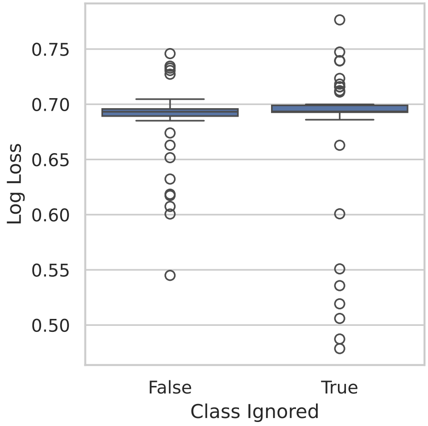


Figure 39: Token class ignored vs. test set loss

While median performance was similar, almost all trials with test set loss below 0.6 ignored token classes (Fig. 39).

**Parameter Initialisation** We test two approaches to weight initialisation, the default PyTorch initialisation, and an approach based on GPT2's weight initialisation [53]. Using the GPT method, we initialise linear models from a normal distribution $\mathcal{N}(0.0, 0.02)$; embeddings are initialised to $\mathcal{N}(0.0, 0.02)$ with zero padding; and layer Normalisation modules are initialised to 1.0 with zero bias. Using PyTorch initialisation, linear layers are initialised from a uniform distribution $U(\frac{-1}{\sqrt{d}}, \frac{1}{\sqrt{d}})$, embeddings are initialised from $\mathcal{N}(0, 1)$, and normalisation layers are initialised to fixed vectors with ones for weights and zeros for biases.

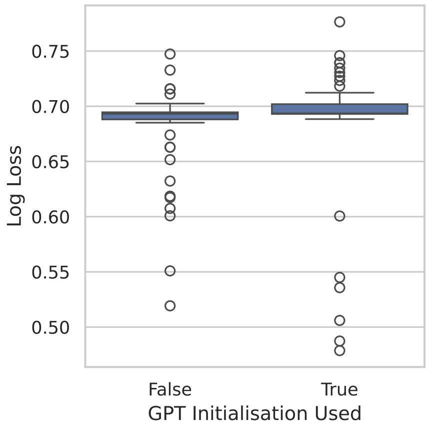


Figure 40: Initialisation method vs. test set loss.

There is no clear advantage to either initialisation function, although the top three trials all used GPT initialisation (Fig. 40).

**Dual Output** We observed during testing that a network with two outputs in the final fully connected layer, optimised using cross entropy, outperformed a network with only a single output using binary cross-entropy (Fig. 41). The dual-output network gives probabilities of gout and non-gout separately, as opposed to only giving a probability of gout. Given an encoder classification-token output $X \in \mathbb{R}^d$, binary cross-entropy loss of a single output using a weight matrix $W \in \mathbb{R}^{1\times d}$, optimises output $\hat{y} = WX$ and loss $l_{bce}$:

$$l_{bce}(\hat{y}, y) = -\left(y \cdot log(\sigma(\hat{y})) + (1-y) \cdot log(1-\sigma(\hat{y}))\right)$$

where $\sigma(\hat{y}) = \frac{1}{1+e^{-\hat{y}}}$.

Dual output cross entropy loss uses $W \in \mathbb{R}^{2\times d}$, and optimises the cross entropy loss $l_{ce}$:

$$l_{ce}(\hat{y}, y) = -\left(log\left(\frac{e^{\hat{y}_1}}{e^{\hat{y}_1}+e^{\hat{y}_2}}\right)y_0 + log\left(\frac{e^{\hat{y}_2}}{e^{\hat{y}_1}+e^{\hat{y}_2}}\right)y_1\right)$$

where $\hat{y} = [\hat{y}_1, \hat{y}_2] = WX$. In this case we say $y_0 = 1$ if and only if $y = 0$ (i.e. the patient does not have gout). Similarly, $y_1 = 1$ if and only if $y = 1$.

Note that these functions are equivalent when $\hat{y}_2 = \hat{y}_1 = \hat{y}$, since $l_{ce}([0, \hat{y}], 1) = l_{bce}(\hat{y}, 1)$ and $l_{ce}([\hat{y}, 0], 0) = l_{bce}(\hat{y}, 0)$. Effectively, the difference is that we use separate weights to learn the true cases and the false cases.

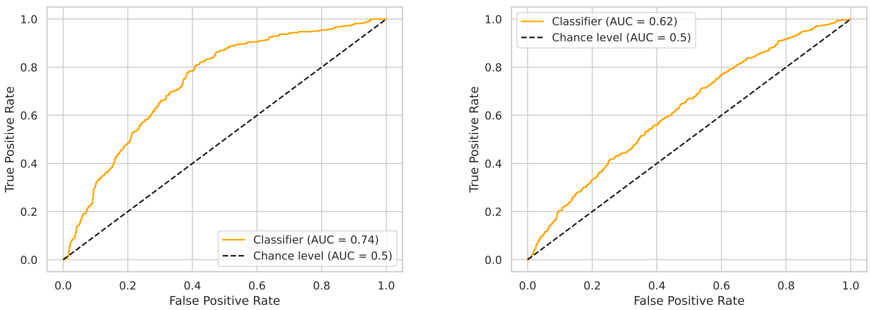


(a) (b)

Figure 41: (a) Dual output layers. (b) Single output layer.

Since these two approaches are not quite equivalent, and can have a significant performance impact, we include both output options in our hyperparameter search. While there is no consistent advantage, the single output model is used in the top three performing trials (Fig. 42).

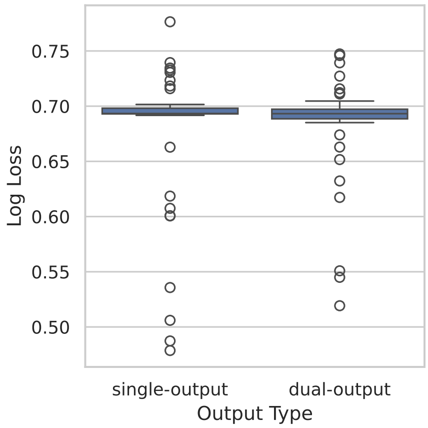


Figure 42: Output type vs. loss.

**Learning Rate** Gradient-descent optimisation methods, including variants like Adam [28] use a learning rate parameter in training to limit the rate at which parameters are changed. Setting the rate too high will result in large changes, potentially overshooting a local minimum and slowing or preventing optimal convergence. Too low a learning rate will slow learning enough that local optima are never reached. We tune the rate by randomly sample learning rates from $log(U(10^{-9}, 10^{-2}))$, where $U$ is the uniform distribution, recording test set loss for each trial (shown in Fig. 43).

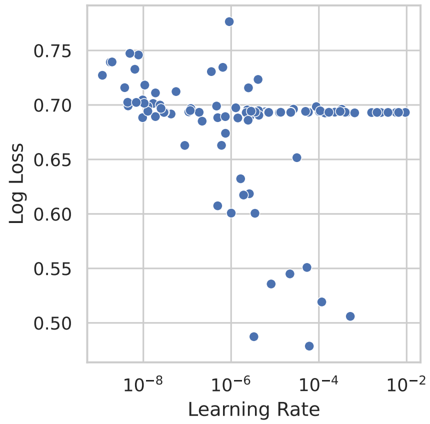


Figure 43: Learning rate vs. test set loss.

While most networks never achieved better test set loss than random (0.693), we see in Fig. 43 that learning rates between $10^{-7}$ and $10^{-}1$ result in the best performance.

#### F.7 Hyperparameter Search Results

For each hyperparameter trial, we record the test set loss at every epoch. These losses are used by the AHSA scheduler [31] to determine which trials to continue and which to halt early. Additionally, they allow us to find the training epoch with the best test set loss, avoiding overfitting. Table 12 summarises the results of the hyperparameter search, including the total running time and the best epoch's test set loss for each trial.

| Trial No. | Iterations | Time (s) | Best Test Loss |
|---|---|---|---|
| 0 | 1 | 621 | 0.730 |
| 1 | 4 | 1590 | 0.693 |
| 2 | 1 | 1144 | 0.693 |
| 3 | 1 | 2740 | 0.693 |
| 4 | 9 | 7018 | 0.618 |
| 5 | 1 | 939 | 0.745 |
| 6 | 2 | 3140 | 0.693 |
| 7 | 1 | 428 | 0.693 |
| 8 | 1 | 957 | 0.693 |
| 9 | 6 | 4787 | 0.632 |
| 10 | 15 | 7252 | 0.607 |
| 11 | 9 | 5541 | 0.600 |
| 12 | 35 | 9491 | 0.600 |
| 13 | 1 | 3420 | 0.689 |
| 14 | 1 | 2599 | 0.701 |
| 15 | 1 | 1573 | 0.698 |
| 16 | 1 | 233 | 0.699 |
| 17 | 1 | 755 | 0.695 |
| 18 | 44 | 29135 | 0.692 |
| 19 | 1 | 1203 | 0.696 |
| 20 | 3 | 4075 | 0.689 |
| 21 | 1 | 1421 | 0.691 |
| 22 | 1 | 245 | 0.723 |
| 23 | 1 | 1474 | 0.693 |
| 24 | 1 | 1792 | 0.696 |
| 25 | 1 | 3022 | 0.693 |
| **26** | **12** | **6160** | **0.478** |
| 27 | 2 | 1693 | 0.693 |
| 28 | 1 | 178 | 0.699 |
| 29 | 5 | 2567 | 0.662 |
| 30 | 7 | 3430 | 0.688 |
| 31 | 2 | 1963 | 0.689 |
| 32 | 4 | 6058 | 0.693 |
| 33 | 1 | 2024 | 0.692 |
| 34 | 1 | 1092 | 0.698 |
| 35 | 2 | 1978 | 0.693 |
| 36 | 1 | 498 | 0.715 |
| 37 | 1 | 1245 | 0.693 |
| 38 | 1 | 507 | 0.693 |
| 39 | 1 | 1550 | 0.693 |
| 40 | 1 | 534 | 0.702 |
| 41 | 1 | 2589 | 0.776 |
| 42 | 1 | 492 | 0.711 |
| 43 | 1 | 723 | 0.696 |
| 44 | 1 | 157 | 0.698 |
| 45 | 1 | 1242 | 0.694 |
| 46 | 1 | 2394 | 0.732 |
| 47 | 7 | 6517 | 0.693 |
| 48 | 14 | 3492 | 0.617 |
| 49 | 4 | 2286 | 0.693 |

| Trial No. | Iterations | Time (s) | Best Test Loss |
|---|---|---|---|
| 50 | 4 | 2982 | 0.550 |
| 51 | 4 | 11151 | 0.673 |
| 52 | 15 | 10890 | 0.688 |
| 53 | 1 | 2141 | 0.694 |
| 54 | 1 | 2239 | 0.734 |
| 55 | 1 | 1576 | 0.693 |
| 56 | 1 | 798 | 0.704 |
| 57 | 4 | 1095 | 0.693 |
| 58 | 15 | 5929 | 0.685 |
| 59 | 1 | 668 | 0.693 |
| 60 | 2 | 4679 | 0.694 |
| 61 | 3 | 1553 | 0.505 |
| 62 | 2 | 1721 | 0.692 |
| 63 | 9 | 2846 | 0.519 |
| 64 | 7 | 4523 | 0.690 |
| 65 | 1 | 475 | 0.718 |
| 66 | 1 | 277 | 0.727 |
| 67 | 1 | 363 | 0.747 |
| 68 | 1 | 1709 | 0.693 |
| 69 | 1 | 3794 | 0.739 |
| 70 | 1 | 847 | 0.692 |
| 71 | 1 | 809 | 0.693 |
| 72 | 7 | 3240 | 0.651 |
| 73 | 1 | 1696 | 0.697 |
| 74 | 1 | 671 | 0.701 |
| 75 | 1 | 795 | 0.693 |
| 76 | 4 | 4571 | 0.693 |
| 77 | 24 | 21380 | 0.662 |
| 78 | 1 | 1829 | 0.694 |
| 79 | 3 | 1420 | 0.702 |
| 80 | 2 | 1082 | 0.693 |
| 81 | 1 | 242 | 0.693 |

The parameters of the best performing trial (no. 26) are shown in Table 13.

| Parameter | Value |
|---|---|
| Training Epochs | 12 |
| Batch Size | 4 |
| Chromosome Embedding Size | 32 |
| Dropout | 0.128 |
| Total Embedding Size | 56 |
| Feedforward Network Internal Scale | 2 |
| Use Gene Embedding Graph | True |
| Gene Embedding Size | 8 |
| Ignore Chromosome | False |
| Ignore Class | True |
| Linformer K | 32 |
| Learning Rate | $6.051 \times 10^{-5}$ |
| Output Layer Type | Single Output |
| Number of Attention Heads | 4 |
| Number of Attention Layers | 3 |
| Position Combination Method | Addition |
| Position Embed Size | 56 |
| Position Encoding | Embedding |
| SNV Embed Size | 16 |
| SNV Encoding | Embedding |
| Use GPT Initialisation | True |

Table 13: Parameters of best-performing trial on test set

This best-performing network has an AUROC of 0.832 in the validation set. Before discussing this further, we first conduct several tests using this architecture that could not practically be included in the hyperparameter search using this network.

#### F.8 Pre-Trained Encoder

Neural networks, and transformers in particular, can use pre-trained to learn underlying patterns and structure of the data. This can significantly improve performance in a number of natural language tasks, particularly when large amounts of related data are available [52]. In pre-training we input partially masked sequences and use the encoder to reconstruct the full sequence, penalising errors. This requires learning relationships between inputs, which will be embedded in the pre-trained network. We then *fine-tune* our pre-trained network to solve the real problem. To determine whether pre-training on the hundreds of thousands of non-gout cases in the UKBB is useful, we compare two variations of the optimal model from Table 13.

In one case, we pre-train an encoder to predict masked SNV tokens using the *LDMed* dataset. In each batch, we mask the classes of 50% of the SNVs at random, then optimise the cross entropy between the output and the true SNV classes, training for two epochs. The encoder is trained using the entire unbalanced training set ($\approx 377,000$ samples) for two epochs, and the training and test set losses are recorded every 100 batches. The SNV class cross entropy loss drops below 0.5 within the first $1,000$ batches, after which it reduces only gradually (Fig. 44).

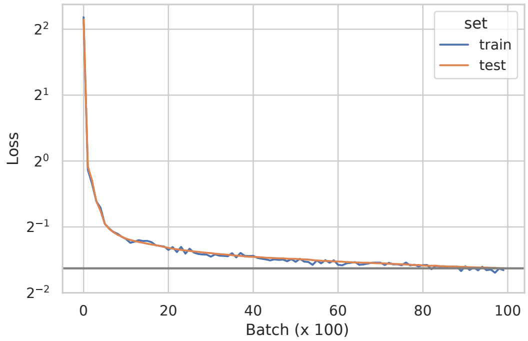


Figure 44: Encoder pre-training loss for every 100th batch on both the training and test sets.

We then train two classifiers, one with random initial weights, and the other using the pre-trained encoder. The randomly initialised version is trained for the optimal number of epochs, 12, according to our hyperparameter search. While the randomly initialised network achieved its optimal test-set loss after 12 epochs, we cannot know whether this will also be true of the pre-trained network. We therefore train for 50 epochs, recording the training and test set losses for each epoch. The per-epoch train and test set errors for both methods are shown in Fig. 45. As we see in Fig. 45, the pre-trained model is able to more easily overfit the data, and as a result does not achieve as low a test error as the randomly initialised model.

While pre-training may be helpful in cases where overfitting is less likely, masked-input pre-training does not improve performance here. It is possible however that a different pre-training objective could bias the network toward a better model of gout, rather than encouraging overfitting.

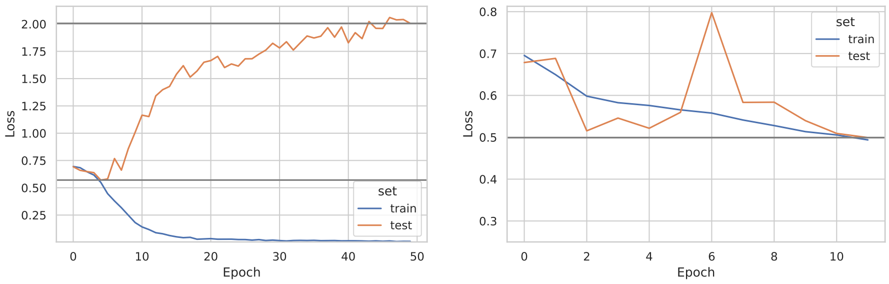


(a) (b)

Figure 45: Training and test set losses vs. number of epochs for pre-trained (a) and randomly initialised (b) transformer-encoder based gout classifiers.

#### F.9 Combined Gout-Urate Training Target

Given the relationship between serum urate levels and gout, it is possible that training to predict urate might help with gout prediction. We investigate this by preparing a model that simultaneously predicts gout and urate, and comparing it to an equivalent model that only predicts gout.

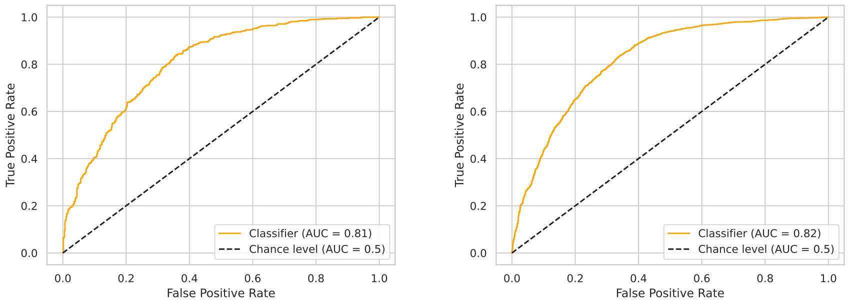


(a) (b)

Figure 46: (a) Combined urate model gout predictions. (a) Plain model gout predictions.

As we see in Fig. 46, there is no significant improvement when adding urate prediction to training. Note that the results in Fig. 46b are from a repeated run with the best-performing hyperparameters, and results vary slightly.

#### F.10 Overfitting and Data Augmentation

Larger models consistently overfit the training data, achieving near perfect accuracy on the training set while failing to generalise to the test set. Data augmentation could mitigate this problem by reducing the number of ways the network can overfit the data. Data augmentation typically involves generating new data using slight variations of existing data that are certain, or at least likely, to have the same correct classification [60]. Generally, data augmentation can be seen as a regularisation technique that encourages decision boundaries further away from training samples, by introducing new training samples in the neighbourhood of existing ones that must share their classification [60].

To test this, we implemented a simple data augmentation method. To increase the training data set from size $n$ to size $n+k$, we include an additional $\frac{k}{2}$ non-gout cases directly from the UKBB data (most non-gout cases are unused to maintain a balanced training set). We randomly draw $\frac{k}{2}$ samples from the existing gout cases to increase the number of gout cases, then in each sample we mask a random continuous chunk of 15% of the SNVs, and replace another 15% of SNVs with random tokens. All phenotypes, including gout, are left the same. Note that since most SNVs remain unchanged, this is most likely still a genome with a high risk of gout. We train a model with the hyperparameters chosen in Appendix F.7 for 18 epochs, and compare this (Fig. 47a) to the results training without data augmentation (Fig. 47b).

Interestingly, overfitting occurs almost immediately with data augmentation enabled, whereas it does not occur until after five epochs without. The best test set loss is nearly identical in both cases, around 0.51.

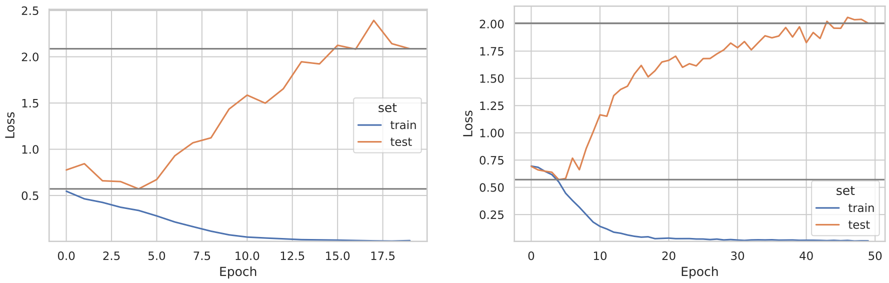


(a)

(b)

Figure 47: (a) Train/test loss during training for augmented model. (b) Train/test loss without data augmentation.
